# Wildlife trade drives animal-to-human pathogen transmission over 40 years

**DOI:** 10.1101/2025.02.09.637309

**Authors:** Jérôme M. W. Gippet, Colin J. Carlson, Tristan Klaftenberger, Mattéo Schweizer, Cleo Bertelsmeier

## Abstract

Wildlife trade affects a quarter of terrestrial vertebrates(*1*) and creates novel opportunities for cross-species pathogen transmission(*2–4*). Yet, its precise role in shaping human-wildlife pathogen exchanges remains unclear. Analysing 40 years of global wildlife trade data, we show that traded mammals are twice as likely to share pathogens with humans than non-traded species. Moreover, time spent in trade strongly predicts the number of zoonotic pathogens that wildlife species host: on average, a species shares an additional pathogen with humans for every 9 years of presence in trade. This work highlights the importance of the wildlife trade in driving animal-to-human pathogen transmission, offering key insights into pathogen biogeography in the Anthropocene and the prevention of future pandemics.

## Main text

Interactions between humans and wildlife promote the transmission of parasites and pathogens(*4*, *5*), which can lead to infectious disease outbreaks(*6*, *7*), resulting in dramatic death tolls and long-lasting socio-economic impacts(*5*, *8*, *9*). Understanding what shapes the spread of zoonotic (i.e., human-transmissible) pathogens and parasites (hereafter pathogens) across species is therefore not only one of the overarching goals of disease ecology, but also a public health priority(*5*, *10*).

The wildlife trade creates opportunities for wildlife-to-human pathogen spillover at each stage of the supply chain, including harvesting, transport, breeding, retail, consumption and companionship(*2*, *5*, *11–14*). For instance, a person buying three variable squirrels in Laotian wildlife markets has been estimated to have an 83% chance of getting at least one leptospirosis-infected individual(*13*). The wildlife trade is therefore often the source of disease outbreaks in humans (*2*, *15*), perhaps most famously including the COVID-19 pandemic, which has been traced back to live mammals sold in the Huanan Seafood Wholesale Market in Wuhan(*16*). While the hunting and consumption of wild meat has been linked to some major epidemics, such as the HIV pandemic(*17*) and some Ebola outbreaks(*18*), other types of wildlife use and trade are also responsible for human infectious disease outbreaks(*2*, *15*). For example, the trade in exotic pets has been implicated in a monkeypox outbreak in 2003 linked to prairie dogs in North America(*19*), and recent hospitalizations due to a rare *Salmonella* strain traced back to pet bearded dragons(*20*).

Despite growing awareness of the epidemic risk posed by the wildlife trade, we still know little about how it affects the exchange of pathogens between animals and humans(*13*, *21–25*). Recent research on the determinants of host-pathogen interactions have mostly focused on environmental, life history, ecological and evolutionary drivers(*7*, *11*, *26–33*). However, the role of human-wildlife interactions associated with the wildlife trade remains poorly studied(*5*, *34*, *35*) and previous empirical studies have only focused on quantifying the number of zoonotic pathogens occurring in traded animals(*22*, *24*, *36*, *37*). As a result, although traded species are known to frequently host zoonotic pathogens(*13*, *21*, *24*), it remains unclear whether species occurring in the wildlife trade are more likely to share pathogens with humans compared to non-traded species. In theory, traded wildlife should be more likely to share pathogens with humans because frequent and close contact with humans increase opportunities for pathogen transmission(*4*, *5*). This can occur either through spillover, where pathogens from wildlife infect humans(*38*) (thus becoming zoonotic), or through reverse zoonosis, where human pathogens (or wildlife pathogens that spilled over humans) infect wildlife (e.g., farmed minks contracting SARS-CoV-2)(*4*, *39*, *40*). Moreover, the more frequently species are traded, the more pathogens they should share with humans.

Here, we tested whether traded wildlife species are more likely to share pathogens with humans compared to non-traded wildlife, and whether the number of years that a species spends on the global wildlife market (in the last 40 years) predicts the number of pathogens it shares with humans. We used the CITES(*41*) and LEMIS(*42*, *43*) trade databases to assess the occurrence of mammal species in the global wildlife trade (Supplementary Fig. 1). The CITES trade database is the most comprehensive database on the international wildlife trade in live animals and animal products. It is global in scale and span over five decades (1975 to present), although, it only contains records for CITES-listed species (∼13% of all mammal species)(*41*, *44*). The LEMIS trade database compiles records of live animals and animal products imported in the United States between 2000 and 2014(*42*, *43*). Although this database is limited to U.S. importations and covers a shorter timespan than the CITES trade database, it includes all species and thus, provides valuable information on the trade in species not recorded by CITES. Here, we focused on the legal trade and excluded all trade records labelled as illegal from both databases, because, by definition, they are not systematically detected and recorded(*45*). In a second analysis, we analysed over 170,000 international trade records in CITES-listed mammals, covering 40 years of legal global trade in wild animals, alive and as products. We focused our analysis on mammals because they are the main reservoir of zoonotic pathogens(*21*, *31*) and the best documented vertebrate clade(*34*, *46*, *47*). We assessed which wild mammal species were known to share pathogens with humans by querying the largest-to-date hand-curated mammal-pathogen associations database(*48*) (>190,000 documented associations between mammals and viral, bacterial, fungal, helminth, and protozoan pathogens and parasites). We accounted for potential biases and confounding factors in all our analyses. Because human-wildlife pathogen exchanges are facilitated by species phylogenetic relatedness with humans(*7*), we accounted for mammals’ phylogenetic distance to humans. Moreover, because pathogens are unequally distributed geographically(*10*, *49*, *50*) and among taxa(*31*, *51*) and because they are also more likely to be detected in well studied species(*29*, *52*), we controlled for taxonomic relatedness, (order, family and genus as nested random effects on the intercept), biogeography (biogeographic realm of origin as random effect on the intercept), and research effort (research effort index as fixed effect; see Methods). Finally, we also accounted for the potential confounding effect of synanthropy on the link between zoonotic pathogens and the wildlife trade. This effect could exist because synanthropic species, that are species that live in or near human-modified environments, are more likely to share pathogens with humans (because of frequent contacts with humans or domesticates(*11*, *29*, *30*, *53*)) and to occur in the wildlife trade (because they are easier to harvest than species occurring only in remote areas(*15*, *54*)).

## Effect of wildlife trade and synanthropy

Among 1,505 traded mammal species, half shared at least one pathogen with humans, while it is the case for only eight percent of non-traded mammals (Fig. 1abc). Using a binomial mixed-effects model, we show that the probability that a mammal species shares pathogens with humans (i.e., risk ratio) is 2.4 times higher in traded than in non-traded species (Odds Ratio [CI95%]_trade_ = 2.72 [2.06–3.6], *p*_trade_ < 0.0001; Fig. 1c) and 1.7 times higher in synanthropic than in non-synanthropic species (Odds Ratio [CI95%]_synanthropy_ = 1.94 [1.43–2.62], *p*_synanthropy_ < 0.0001; Fig. 1c; see details in Supplementary Text 1). We then used a structural equation model (SEM) to test whether the probability of sharing pathogens with humans is directly linked to trade, or the result of a confounding effect of synanthropy, while accounting for the links between these variables and research effort (Fig. 1d). Our SEM supports a direct positive effect of trade on the probability that wild mammals share pathogens with humans (Std. est. = 0.14, R^2^_marginal_ = 0.49; Fig. 1d; Supplementary Text 2). In addition, synanthropy was positively associated with both zoonotic host status (Std. est. = 0.08) and trade status (Std. est. = 0.15). However, our SEM shows limited explanatory power for the effect of synanthropy on trade (R^2^_marginal_ = 0.02; Fig. 1d), and a weak effect of synanthropy on zoonotic host status (compared to other effects in the SEM; Fig. 1d). This indicates that synanthropy does not act as a confounding factor on the link between trade and human-wildlife pathogen sharing. Interestingly, our SEM supports a positive effect of trade and synanthropy on research effort, thus suggesting that part of the positive link between research effort on zoonotic host status is mediated by these two variables (i.e., traded and synanthropic species are more studied and thus their pathogens are better known; Fig. 1d). Overall, these results support our main hypothesis that, even after accounting for sampling bias, species that are present in the wildlife trade are more likely to share pathogens with humans, because repeated and close cross-species contact creates opportunities for pathogen transmission. However, while some species have been continuously traded in the last four decades, others only sporadically occur in the global wildlife trade(*55*, *56*) (Fig. 2a). Therefore, the opportunities for cross-species contact and thus pathogen transmission may strongly differ from one species to the other.

**Fig. 1.**
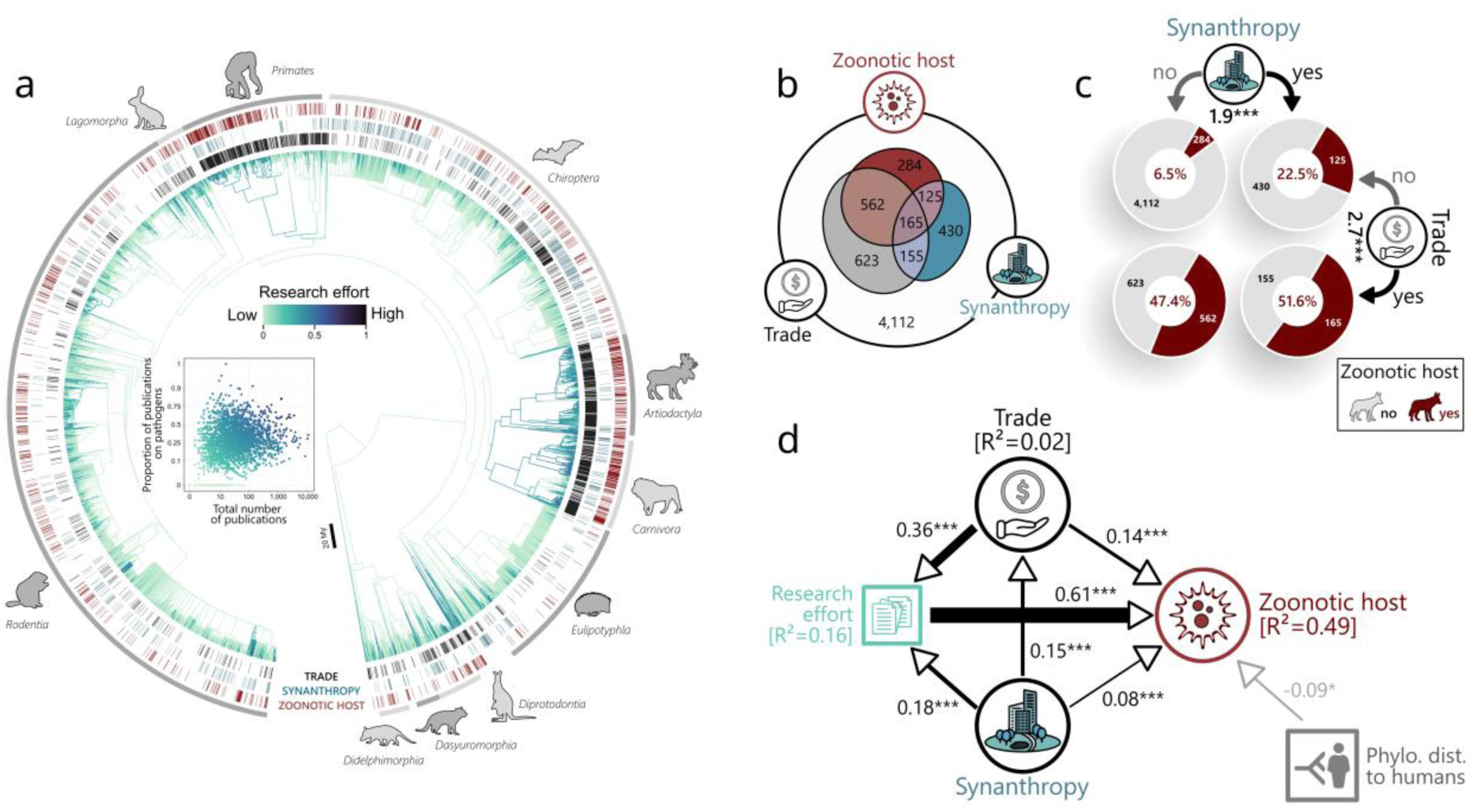
The probability of sharing pathogens with humans is higher in traded and synanthropic species. **a**, Distribution of traded, synanthropic, and zoonotic host mammals (i.e., species known to share at least one pathogen with humans) among extant wild mammal species (5,565 of the 6,471 species shown in the phylogram). The colour of the branches represents the research effort index used to account for knowledge bias in statistical analyses. The ten species richest mammal orders are indicated (silhouettes from PhyloPic(*57*)). **b,** Overlap between traded, synanthropic, and zoonotic host species. **c,** Proportions of zoonotic host species (in red) among traded and synanthropic mammals. The number of species in each group is also indicated. Numbers next to icons are standardized effect sizes (odds ratios) of trade and synanthropy on the probability of sharing pathogens with humans (***: p<0.001). **d,** Structural equation model (SEM) showing the strength and significance of hypothesised causal relationships between trade, synanthropy, research effort, phylogenetic distance to humans and zoonotic host status among wild mammals. Black arrows indicate positive links between variables and grey arrows negative ones. Numbers along arrows are standardized SEM estimates (*: p>0.05, ***: p<0.001). Arrows’ width is proportional to these estimates.

**Fig. 2.**
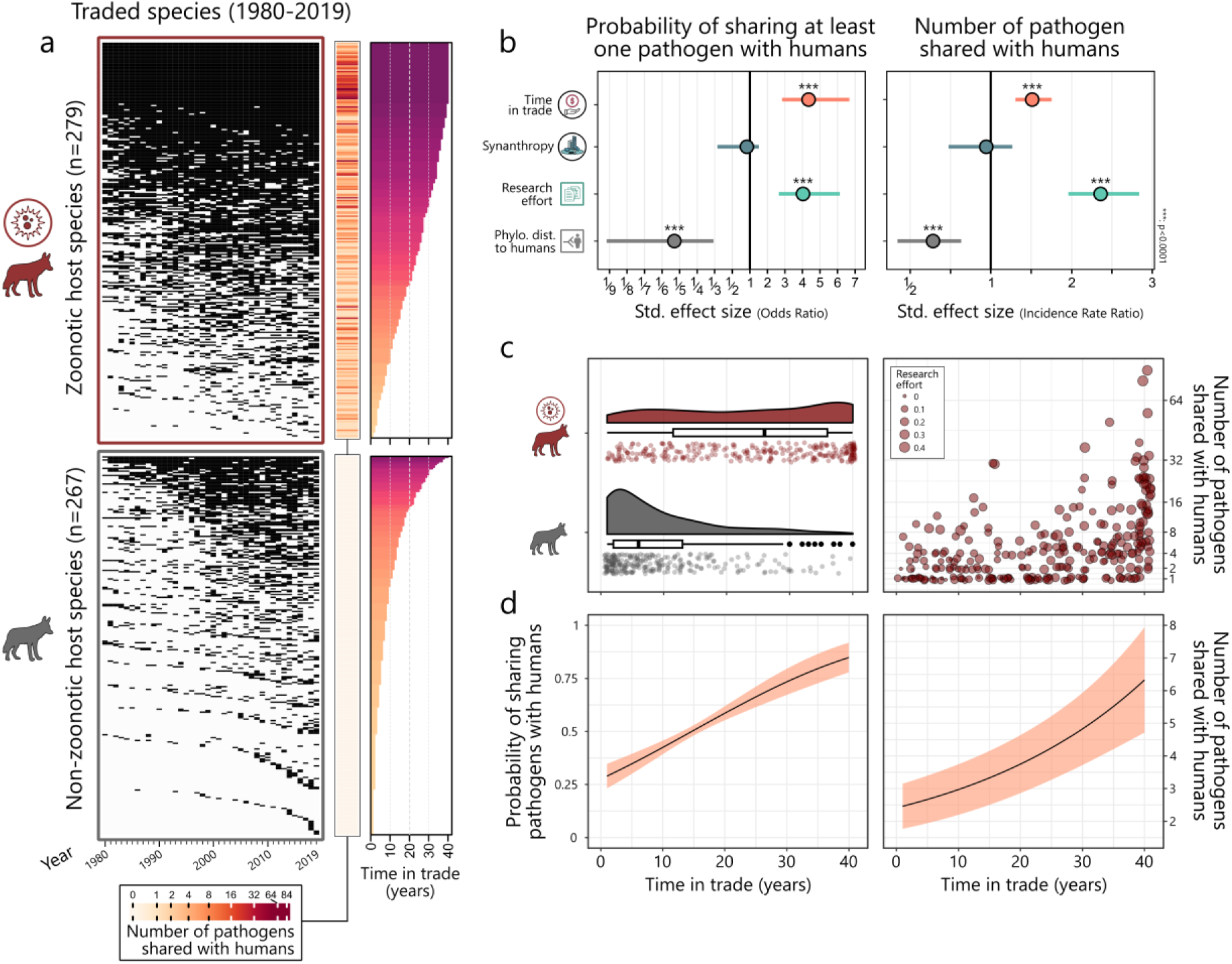
Frequently traded species are more likely to share pathogens with humans. a, On the left, barcode plot representing mammal species occurrence in trade in the last 40 years. Each line is a mammal species that has been traded at least once between 1980 and 2019 (n = 546), each column is a year (black cells indicate trade). Lines (i.e., species) are ordered by zoonotic host status and time in trade (and time since first trade event). **b,** Standardized effect sizes (estimate ± 95% CI) of predictors (fixed effects) in the two sub-models of the hurdle negative binomial mixed effect model (zero-inflation sub-model on the left, conditional sub-model on the right). The x-axis has been deliberately modified to allow an intuitive and undistorted comparison of positive and negative effects. **c,** On the left, time in trade among species sharing pathogens with humans or not. On the right, number of pathogens shared with humans as function of time in trade. Each point is a mammal species. The size of points is proportional to research effort. The vertical axis is square rooted to improve readability. **d**, Average effects (slope ± CI95%) of time in trade on the probability of sharing at least one pathogen with humans (left) and on pathogen richness (right).

## Time in trade predicts zoonotic pathogen richness in wildlife

Species that have been traded more frequently and for longer periods of time have had more opportunities to exchange pathogens with humans and should therefore be more likely to host a greater number of zoonotic pathogens. To test this, we performed a temporal analysis of the global trade in wild mammals over the last 40 years (1980-2019) using the CITES trade database, the only long-term longitudinal dataset of the global wildlife trade(*41*). We analysed 171,749 trade records in 546 mammal species that have been traded at least once between 1980 and 2019 (among species listed in CITES Appendices I or II; see Methods) and calculated the frequency at which species were traded in the last 40 years, as the number of years with recorded trade (hereafter “time in trade’; Fig. 2a). Using a hurdle negative binomial mixed-effect model (Supplementary Text 3-4 for detailed results), we show that, as expected, time in trade increases both the probability of sharing at least one pathogen with humans (Zero-inflation sub-model: Odds Ratio [CI95%] = 4.4 [2.8–6.7], *p* < 0.0001; Fig 2b,c,d) and the number of zoonotic pathogens that a mammal species hosts (Conditional sub-model: Incidence Rate Ratio [CI95%] = 1.5 [1.3–1.8], *p* < 0.0001; Fig 2b,c,d). Using this model, we estimate that, on average over the period considered, a wild mammal species shares one additional pathogen with humans every 9 years of presence in trade (mean [CI95%] = 8.9 [7.4 – 11.2], see Supplementary Text 5 for calculation details). This implies that pathogens hosted by traded species that currently do not infect humans are likely to do so in the future. These results highlight the dynamic nature of human-wildlife pathogen interaction networks and parallel the positive correlation observed between time since domestication and number of pathogens shared with humans in domesticated mammals(*58*). Although these two trends are difficult to compare (in part because animal domestication is much older than the global wildlife trade), they both highlight the importance of sustained and close contact between animals and humans in promoting cross-species pathogen transmission and in shaping the host-pathogen associations that exist today.

Animal-to-human transmission (spillover) is the most likely driver of trade-related pathogen exchange between wild animals and humans, because of the asymmetry of human-animal interactions (e.g., humans regularly consume wildlife and livestock, while the opposite is extremely rare)(*4*), and because wildlife frequently acts as a reservoir for human-transmissible pathogens, while the opposite is less common(*59*). Understanding the precise processes responsible for the patterns highlighted in this study will require more detailed data on the temporal sequence of pathogen transmission across animals and humans(*32*). Genomic tools will be central to retrace pathogen transmission across species(*17*, *60–63*) and interfaces (e.g., *Mycobacterium bovis* transmission across wildlife and livestock in South Africa(*64*)), and will accelerate the compilation of detailed records of pathogen prevalence in human and animal populations (wild, captive and domesticated(*63*, *65–67*)). Moreover, museomics (the study of genomic data obtained from ancient and historic DNA from specimens in museum collections(*68*)) now allows to identify and quantify the prevalence of pathogens that infected wild and captive animals decades to centuries ago(*69–71*). Our analysis also uses on a simplified representation of host-pathogen interactions (i.e., presence/absence) and thus overlook both inter- and intra-specific variation in pathogen prevalence(*63*, *65*). Higher resolution pathogen testing data will be crucial in assessing whether traded species are reservoirs with a high risk of pathogen transmission, common but dead-end hosts with low transmission risk, or anecdotal hosts(*72*). This information is essential to accurately identify high-risk species and focus regulations on them, rather than using total trade bans that could dilute control efforts or even backfire if trade is massively diverted to illegal channels(*73*).

## Conclusions

Overall, our findings underscore the urgent need to enhance efforts to screen traded animals and animal products for pathogens and evaluate their potential for transmission to humans(*12*, *13*). As this study focuses on species traded legally at international scale, it overlooks local but widespread wildlife markets (e.g., wild meat markets(*74*, *75*) or local exotic pet markets(*76*)), as well as illegal markets(*22*). Accounting for these components of the wildlife trade might improve our ability to evaluate risks related to cross-species transmission, and ultimately, to better predict and prevent future zoonotic disease outbreaks. However, it will require important international efforts to continuously survey and compile information on these markets (e.g., identity and abundance of traded species, purpose of trade, presence and intraspecific prevalence of pathogens). Currently, this information is rare and scattered among numerous independent studies with an uneven geographical and temporal representation(*13*, *75*, *77*, *78*).

The COVID-19 pandemic sparked a thorough re-evaluation of current wildlife trade regulations, exposing critical gaps in our ability to monitor and limit disease outbreaks linked to exploited and traded species(*79*). Currently, the primary international agreement regulating the wildlife trade, CITES, focuses exclusively on preventing species extinction caused by overexploitation of natural populations(*80*). Our findings are thus critical for ensuring the effectiveness of upcoming regulations aimed at pandemic prevention, including potential reforms to CITES(*81*), the WOAH/CITES collaborative agreement(*82*) and the proposed WHO pandemic agreement(*83*). By showing that traded wildlife species share one additional pathogen with humans for every decade of presence in the global wildlife market, our findings underscore that mitigating future zoonotic pathogen emergence will require reducing the volume of wildlife trade – including both species that pose a current risk to human health, and those that might someday soon, given the opportunity.

## Acknowledgments

We thank O. Bates for suggestions that improved previous version of this work.

## Fundings

Canton de Vaud, Switzerland (CB)

Swiss National Science Foundation SNSF, grant 310030_192619 (CB)

SERI-funded ERC grant SPREAD, MB22.00086 (CB)

U.S. National Science Foundation, NSF DBI 2515340 (CJC)

## Author contribution

Conceptualization: JMWG, CJC, CB

Data curation: JMWG, CJC, MS

Formal analysis: JMWG, TK

Investigation: JMWG

Supervision: JMWG, CB

Validation: JMWG, CJC

Visualization: JMWG

Writing – original draft: JMWG

Writing – review & editing: JMWG, CC, TK, MS, CB

## Conflict of interests

Authors declare that they have no competing interests.

## Data and materials availability

All data and code used in the study are available at

https://github.com/JGippet/WildlifeTrade_ZoonoticPathogens [repository will be made public upon publication].

## List of supplementary Materials

Materials and methods

Supplementary Materials (includes figures, tables and text)

## Materials and methods

### Taxonomic harmonization

We combined multiple databases on mammals’ taxonomy, biogeography, zoonotic pathogens, and trade (Supplementary Table 1). To assemble these databases, we performed a taxonomic harmonization procedure using, as a reference, the Mammals Diversity Database (MDD), which is the most complete and up-to-date authoritative database on mammal taxonomy(*1*). For each database to collate, we first matched species names against MDD’s accepted names. Then, for species that were not found in MDD, we matched them against all possible synonyms of MDD accepted names using the R package *taxize*(*2*). Species that could not be linked to a MDD species after this step were excluded from further analyses (depending on the dataset, 93-99% of species were matched; Supplementary Table 2). Because sub-species over-complexify cross-referencing and are subject to frequent reclassifications and disagreements among taxonomists(*3*), we chose to merge them at the species level. Our final dataset listed 6,456 valid mammal species (excluding species listed as extinct (99 species) and domesticated (38 species, including wild relatives) in MDD, and *Homo sapiens*; Supplementary Table 3). Taxonomic classification (order, family, genus) and the biogeographical origin (Afrotropic, Australasia/Oceania, Indomalaya, Marine, Nearctic, Neotropic, Palearctic) of each species were also extracted from the MDD database.

### Host–parasite association data

To assess whether mammals are known to host pathogens and parasites (hereafter ‘pathogens’) that are also known to infect humans, we used the CLOVER database(*4*) (downloaded on February 29^th^, 2023), a reconciled aggregate of four extensive host-pathogen associations databases: GMPD 2.0 (Global Mammal Parasite Database(*5*)), EID2 (ENHanCEd Infectious Diseases Database(*6*)), HP3 (Host-Parasite Phylogeny Project(*7*)), and Shaw database(*8*). The CLOVER database has been used in previous studies to assess the occurrence of zoonotic pathogens in mammals(*9–11*) and is the highest quality host-pathogen association dataset to date, as it has been manually cleaned to remove errors and harmonise taxonomy. After taxonomic harmonization and data filtering (e.g., removing extinct and domesticated species), the dataset contained a total of 14,383 unique associations, including 8,188 involving zoonotic pathogens, between 1,312 mammal species (incl. humans) and 4,446 pathogens (incl. 2,424 zoonotic pathogens), including viruses (708, incl. 400 zoonotic), bacteria (1,752, incl. 1,509 zoonotic), fungi (330, incl. 297 zoonotic), helminths (1,289, incl. 148 zoonotic), and protozoa (367, incl. 70 zoonotic). A total of 1,144 mammal species shares at least one pathogen with humans. Using this information, we calculated the number of zoonotic pathogen species per mammal species.

### Research effort

To account for knowledge bias, we assessed research effort by retrieving the number of publications per mammal species using the *easyPubMed* R package(*12*). We computed two indices of research effort per species by searching for keywords in the title and abstract of publications listed in PubMed(*13*). First, we calculated the total number of publications per mammal species, based on all possible valid specific Latin names and all possible common names (in English) from MDD. Second, we calculated the proportion of all publications that was focused on diseases or pathogens by refining our previous search with the following string: “pathogen* OR zoono* ‘public health’ OR ‘emerging infectious disease’ OR ‘disease’ OR parasit* OR virus OR viral OR helminth* OR bacteria* OR fung* OR protozoa OR infect*”. We finally computed a unique index of research effort per species by multiplying the total number of publications (log-transformed and rescaled to [0-1]) to the proportion of these publications focusing on diseases and pathogens (logit-transformed and rescaled to [0-1]). The equation for calculating this index is as followed:

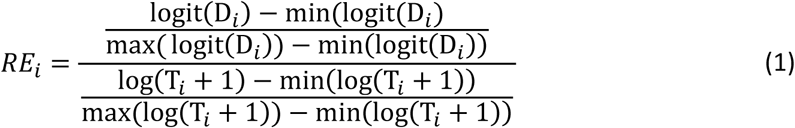

With 𝑅𝐸_𝑖_ the index of research effort for species *i*, 𝐷_𝑖_ the proportion of all publications that focused on diseases or pathogens for species *i* (to avoid infinite values, the logit of 0 and 1 were calculated as the logit of 0.025 and 0.975, respectively), and 𝑇_𝑖_ the total number of publications for species *i*. This index of research effort combines the information on the total number of publications and proportion of these publications focused on pathogens and diseases (see Fig. 1a).

### Phylogenetic distance to humans

We used the VertLife database(*14*) to access phylogenetic relationships between mammals (downloaded on February 29^th^, 2023). We calculated each species average phylogenetic distance to humans (in million years) from 20 randomly selected trees.

### Synanthropy

We assessed mammals’ propensity to live inside or near humans and human settlements using the IUCN Red List database(*15*). A mammal species was listed as synanthropic if it was known to occur in urban areas (habitat code 14.4), rural gardens (14.5), water storage areas (15.1), aquaculture ponds (15.3), wastewater treatment areas (15.6) and canals and drainage channels (15.9). A total of 875 mammal species (out of 6,456) were considered as synanthropic (14%).

### Mammals in the global wildlife trade

We used the CITES trade database(*16*) and the LEMIS trade database(*17*) to assess whether species were traded or not (following recommendations in characterizing wildlife trade(*18*)). The CITES trade database is the most comprehensive database on the international wildlife trade in live animals and animal products and contains records for CITES-listed species from 1975 to present(*16*, *19*). The LEMIS trade database compiles records of wildlife and wildlife products imported in the United States between 2000 and 2014(*17*, *20*). Although this database is limited to U.S. importations and covers a shorter timespan than the CITES trade database, it covers all species and thus, provides valuable information on the trade in species not listed in CITES appendices and therefore absent from the CITES trade database. In this work, we considered species traded legally alive or as products, for commercial, personal, medical, scientific, and hunting trophy purposes. These trade purposes represent the majority of CITES and LEMIS trade records (84% and 99%, respectively). We excluded illegal trade records because they are not systematically reported(*21*) and cannot be easily compared with legal trade records.

### Temporal analysis of the global wildlife trade

The CITES trade database is the only long-term (1975-present) and global compilation of the trade in wild animals(*19*). We used the aggregated version of the CITES trade database(*16*) (Comparative tabulation version 2023.1) to avoid duplicated trade records (e.g. when both importers and exporters report the same trade event). The CITES trade database totals 405,066 records of trade in mammals since 1975 (after taxonomic harmonisation). However, we applied several filters to the database, thus reducing the number of analysed records to 290,241. First, we chose to consider only the period 1980-2019 (40 years) because the first five years (1975-1979) and the last three years (2020-2022) of recording are incomplete, due to non-systematic recording during the first years (many species were not yet listed in CITES and many countries were not yet parties of CITES before 1980(*22*)), and of a time lag in reporting for the recent years(*16*). We also excluded species that were listed in CITES after 1980 because their trade was not reported systematically over the period 1980-2019, and species listed in CITES appendix III because their trade is not systematically recorded at global scale as CITES agreements protect (an report) them only when they are sourced from some specific countries(*23*). Using this refined dataset, we then calculated for each traded mammal species the number of years that the species appeared in trade over the last 40 years.

### Statistical analysis

All data processing, statistical analyses and visualizations were performed in R and Rstudio(*24–28*). Generalised linear models were computed using the *glmmTMB* package(*29*). The statistical validity of each model was systematically assessed using the *performance*(*30*) and *DHARMa*(*31*) packages. For mixed effects models, whenever random effects with an estimation very close to zero created a singularity issue (that can hinder model’s convergence), the problematic factors were removed from the model to allow a non-singular fit (following package recommendations(*32*)).

We used the sjPlot package(*33*) to calculate effect sizes of our GLMs. To interpret effect sizes expressed as odd ratios, we converted them in risk ratios following (eqn. 2). Sequential equation models were computed using the *piecewiseSEM* package(*34*).

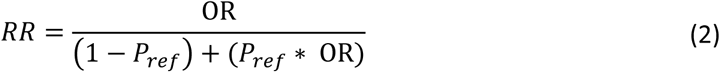

With RR the risk ratio, OR the odd ratio, and 𝑃_𝑟𝑒𝑓_ the baseline prevalence (i.e., the incidence in the baseline group).

### Effect of trade on the probability of sharing pathogens with humans

To assess whether, among all mammal species, traded species are more likely to share pathogens with humans, we used a binomial mixed-effects model (logit link function) with zoonotic host status (i.e., the species shares at least a pathogen with humans or not) as response variable and trade status (i.e., traded or not) as predictor (fixed effect). To account potential confounding effects, we added the index of research effort (see eqn. 1), synanthropic status (i.e., synanthropic or not), and phylogenetic distance to humans as fixed-effect covariates. To account for the taxonomic relatedness between species, we added mammalian order, family and genus as nested random effects on the intercept. Species belonging to genera, families and orders with less than three representatives were excluded to avoid issues in the computation of random effects. To account for the biogeographical origin of mammal species, we added their biogeographical realm as a random effect on the intercept. Species occurring in two or more biogeographical realms were assigned to the category ‘multiple’.

To further test whether the statistical relationship between trade and the probability of sharing pathogens with humans was direct or the result of the confounding effect of synanthropy, we used a structural equation model (SEM)(*34*). SEM is a multivariate analysis technique that integrates multiple predictor and response variables into a single directed network of hypothetical causal links, represented through linear equations. SEM enables the estimation of the strength and significance of each relationship in the model (using standardized estimates) while accounting for all specified relationships within the network(*34*), therefore allowing to account for the potential confounding effect of synanthropy on the relationship between trade and the probability of sharing pathogens with humans. Moreover, this modelling technique also allows to specify networks of interdependence between our variables of interest (zoonotic host status, trade and synanthropy) and research effort. Our SEM model consists in the combination of three mixed effects sub-models, all accounting for the taxonomic relatedness between species (using mammalian order, family and genus as nested random effects on the intercept), and for the biogeographical origin of mammal species (using biogeographical realm as a random effect on the intercept). The first sub-model is a Gaussian model with research effort as the response variable and trade status and synanthropy status as predictors. This sub-model results from the hypothesis that species that are traded and synanthropic are more likely to be studied. The second sub-model is a binomial model (logit link function) with trade status as response and synanthropy as predictor. This sub-model results from the hypothesis that synanthropy increases the probability of being traded. The third sub-model is a binomial model (logit link function) with zoonotic host status as response and trade status, synanthropy status, phylogenetic distance to humans, and research effort as predictors. This sub-model results from the hypotheses that the probability of being known to share zoonotic pathogens with humans increases with phylogenetic proximity to humans, research effort, synanthropy and trade.

### Effect of time in trade on the number of pathogens shared with humans

We tested whether the number of years that a species has occurred in trade is positively linked to the probability of sharing pathogens with humans and to the number of pathogens shared. Here, we focused on mammal species listed in CITES appendices I and II, that were traded at least once between 1980 and 2019, based on records of the CITES trade database(*16*). We also excluded species that were listed in CITES after 1980 because their trade was not reported systematically over the period 1980-2019, and species listed in CITES appendix III because their trade is not systematically recorded at global scale as CITES agreements protect them only when they are sourced from some countries(*23*). Our analysis thus included 546 mammal species for which international trade events were recorded globally and continuously between 1980 and 2019. We then tested whether time in trade is positively linked to the probability of sharing pathogens with humans and to the number of pathogens shared with humans using a hurdle negative binomial mixed-effects model with pathogen richness as response variable (count, between 0 and 84) and time in trade (number of years, between 1 and 40) as predictor (i.e., fixed effects). We also added synanthropy (binary, synanthropic or not), phylogenetic distance to humans and research effort (see eqn. 1) as a fixed-effect covariates. To account for the taxonomic relatedness between species, we added mammalian order, family and genus as nested random effects on the intercept, to account for the biogeographical origin of mammal species, we added their biogeographical realm as a random effect on the intercept. After running the full model (i.e., all effects in both zero-inflation and conditional sub-models), random effects with low variance were removed to avoid singularity issues (Supplementary Text 3).

### Sensitivity analysis

To assess the robustness of our results to variations in the host-pathogen association data used to test our hypotheses, we applied two independent filters to the CLOVER dataset, resulting in four distinct combinations (and thus datasets). We then repeated our analyses on each dataset and compared the outcomes. First, we excluded host-pathogen associations from the EID2 database (one of the four databases compiled in CLOVER), as EID2 includes a substantial number of records from experimental infections(*6*). These records could introduce artificial host-pathogen associations and potentially inflate the number of species sharing pathogens with humans. Second, we filtered out all host-pathogen associations involving bacteria, helminths, fungi, and protozoa, focusing exclusively on viruses. This allowed us to determine whether our findings hold when considering only the pathogen group that poses the highest pandemic risk in the future(*35*). All our results remained consistent across both sources of variation in the host-pathogen association dataset. For a detailed description of the methods and results of this sensitivity analysis, see Supplementary Text 6.

---

**Fig. S1:**
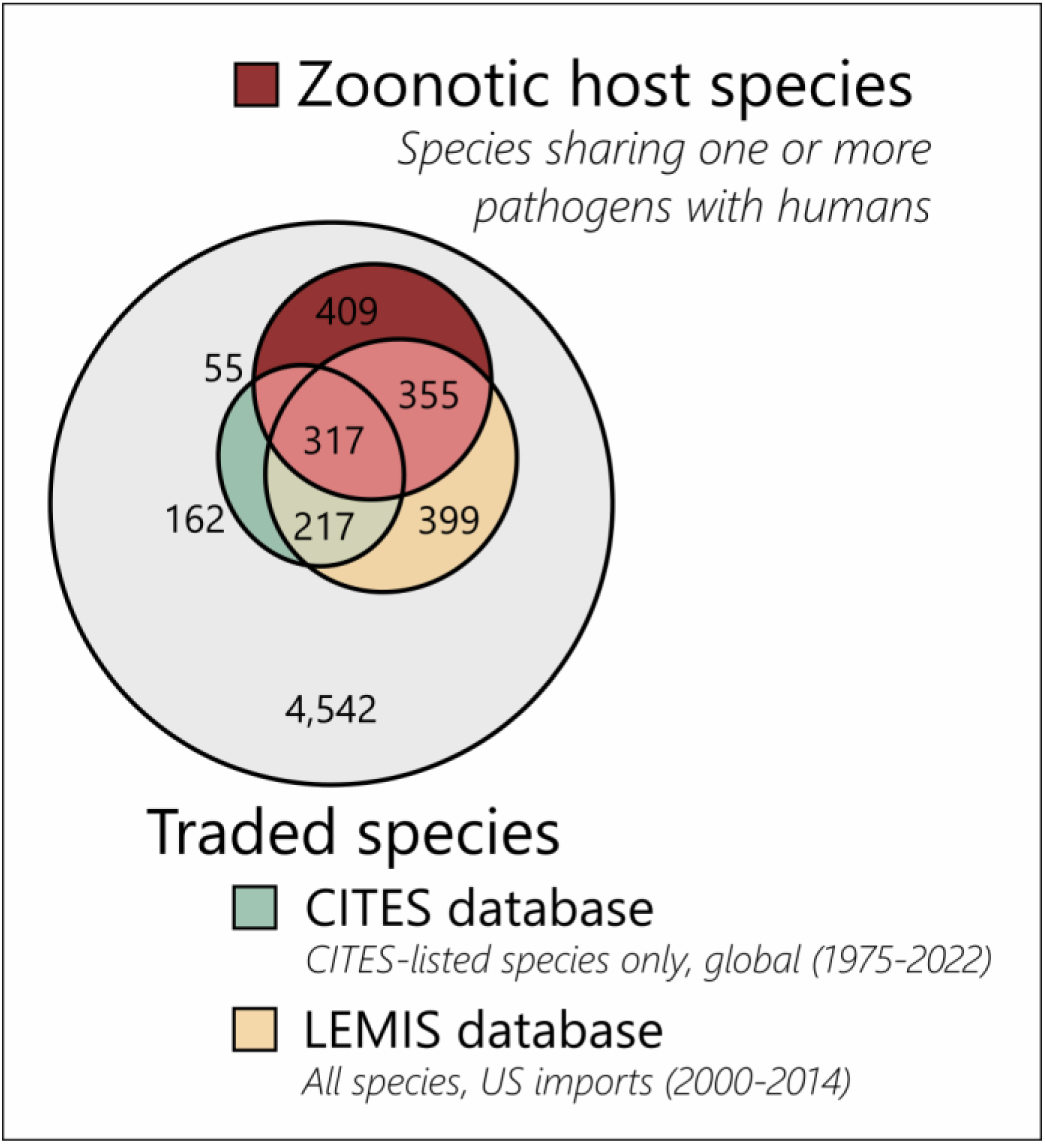
Species overlap between trade databases (CITES and LEMIS). This Euler plot does not include domesticated and extinct species.

**Table S1:** List of databases used in this study.

| Database | Use | Version or access date | Link |
| --- | --- | --- | --- |
| Mammals Diversity Database | Taxonomy (reference database) | v1.9.1 | https://www.mammaldiversity.org/ |
| CLOVER | Host-pathogen interactions | Accessed on February 29th, 2023 | https://github.com/viralemergence/clover |
| CITES checklist | CITES appendices and date of listing | Accessed on June 30th, 2023 | https://checklist.cites.org/#/en |
| CITES trade database | Traded species, Temporal trends in species trade | v2023.1 | https://trade.cites.org/ |
| LEMIS | Traded species | v2020 | https://github.com/ecohealthalliance/lemis |
| IUCN Red list database | Synanthropic species | Accessed on May 1st, 2024 | https://www.iucnredlist.org/ |
| VertLife | Mammal phylogeny | Accessed on February 25th, 2023 | http://vertlife.org/phylosubsets/ |

**Table S2:** Matching rates with the reference database (MDD) for the five main sources of information used in this study.

|  | Initial number of species | Number of species matched to MDD | Matching rate (%) |
| --- | --- | --- | --- |
| CLOVER | 1364 | 1341 | 98 |
| CITES | 838 | 783 | 93 |
| LEMIS | 1449 | 1348 | 93 |
| IUCN (synanthropic species) | 1770 | 1746 | 99 |
| Mammal phylogeny | 5911 | 5658 | 96 |

**Table S3:**
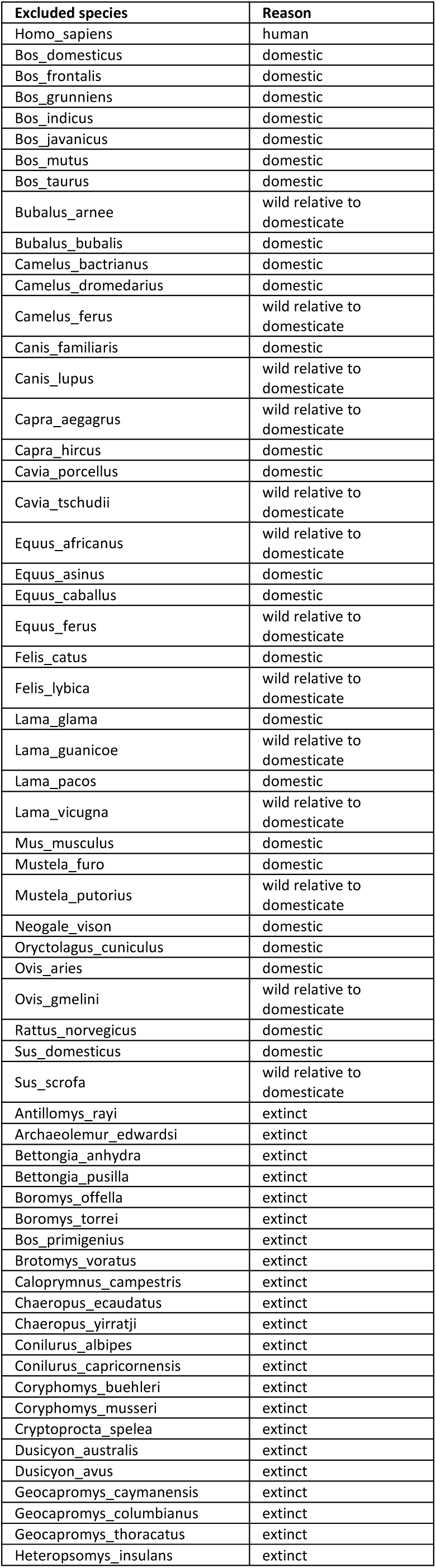
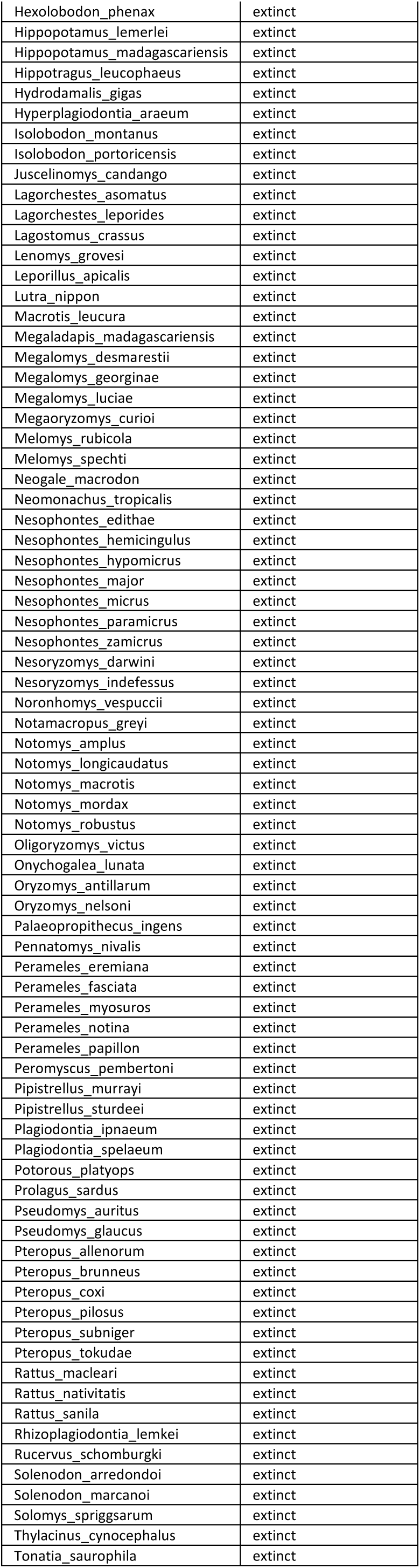
List of mammal species excluded from the study and reason for exclusion.

## Supplementary Text 1

Detailed results for the binomial mixed effects model (presented in Fig. 1c) used to test whether traded species are more likely to share pathogens with humans, while accounting for potential confounding effects.

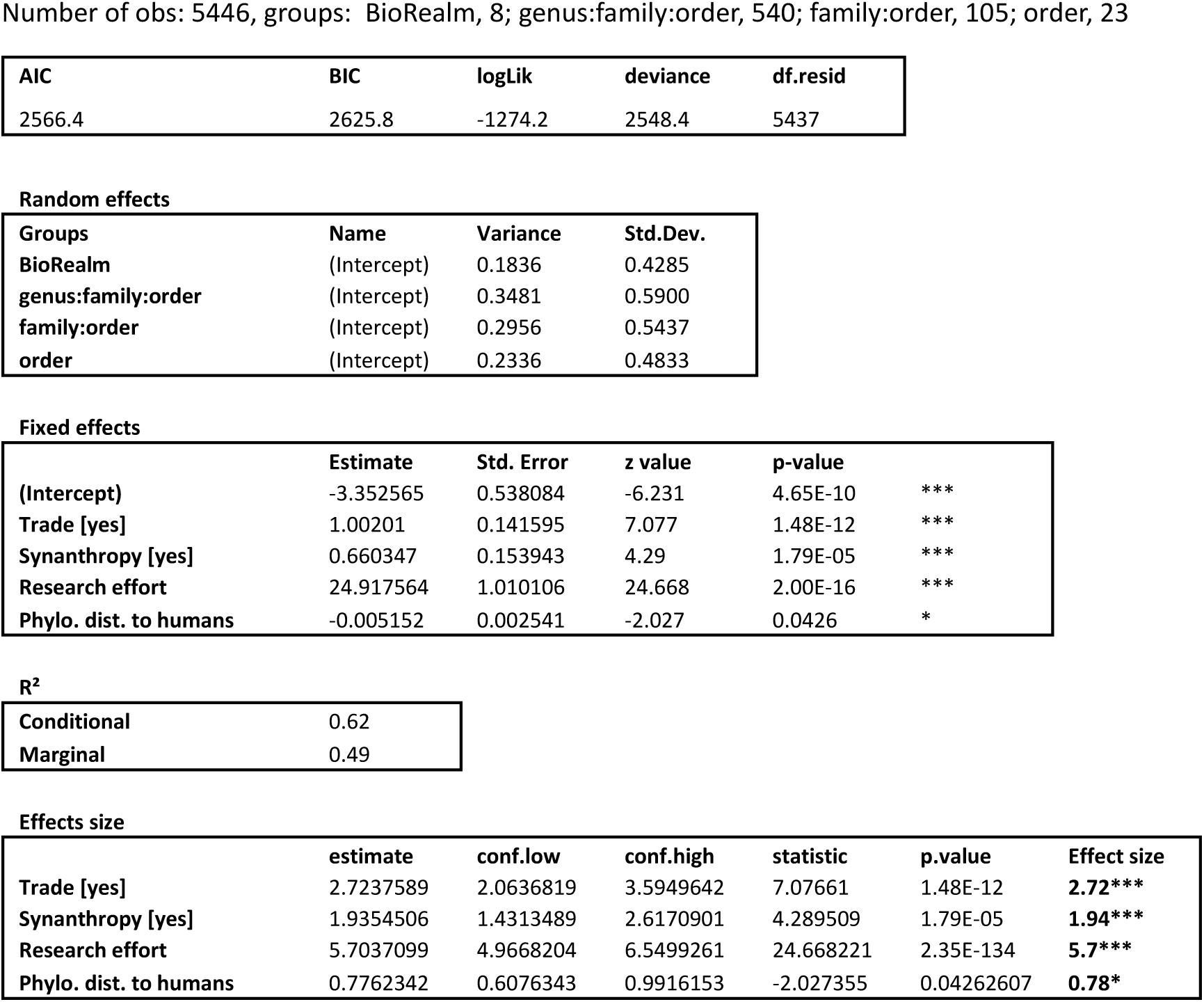

| AIC | BIC | logLik | deviance | df.resid |
| --- | --- | --- | --- | --- |
| 2566.4 | 2625.8 | -1274.2 | 2548.4 | 5437 |

Random effects
| Groups | Name | Variance | Std.Dev. |
| --- | --- | --- | --- |
| BioRealm | (Intercept) | 0.1836 | 0.4285 |
| genus:family:order | (Intercept) | 0.3481 | 0.5900 |
| family:order | (Intercept) | 0.2956 | 0.5437 |
| order | (Intercept) | 0.2336 | 0.4833 |

Fixed effects
|  | Estimate | Std. Error | z value | p-value |  |
| --- | --- | --- | --- | --- | --- |
| (Intercept) | -3.352565 | 0.538084 | -6.231 | 4.65E-10 | *** |
| Trade [yes] | 1.00201 | 0.141595 | 7.077 | 1.48E-12 | *** |
| Synanthropy [yes] | 0.660347 | 0.153943 | 4.29 | 1.79E-05 | *** |
| Research effort | 24.917564 | 1.010106 | 24.668 | 2.00E-16 | *** |
| Phylo. dist. to humans | -0.005152 | 0.002541 | -2.027 | 0.0426 | * |

R<sup>2</sup>
|  |  |
| --- | --- |
| Conditional | 0.62 |
| Marginal | 0.49 |

Effects size
|  | estimate | conf.low | conf.high | statistic | p.value | Effect size |
| --- | --- | --- | --- | --- | --- | --- |
| Trade [yes] | 2.7237589 | 2.0636819 | 3.5949642 | 7.07661 | 1.48E-12 | **2.72\*\*\*** |
| Synanthropy [yes] | 1.9354506 | 1.4313489 | 2.6170901 | 4.289509 | 1.79E-05 | **1.94\*\*\*** |
| Research effort | 5.7037099 | 4.9668204 | 6.5499261 | 24.668221 | 2.35E-134 | **5.7\*\*\*** |
| Phylo. dist. to humans | 0.7762342 | 0.6076343 | 0.9916153 | -2.027355 | 0.04262607 | **0.78\*** |

**Fig. S2:**
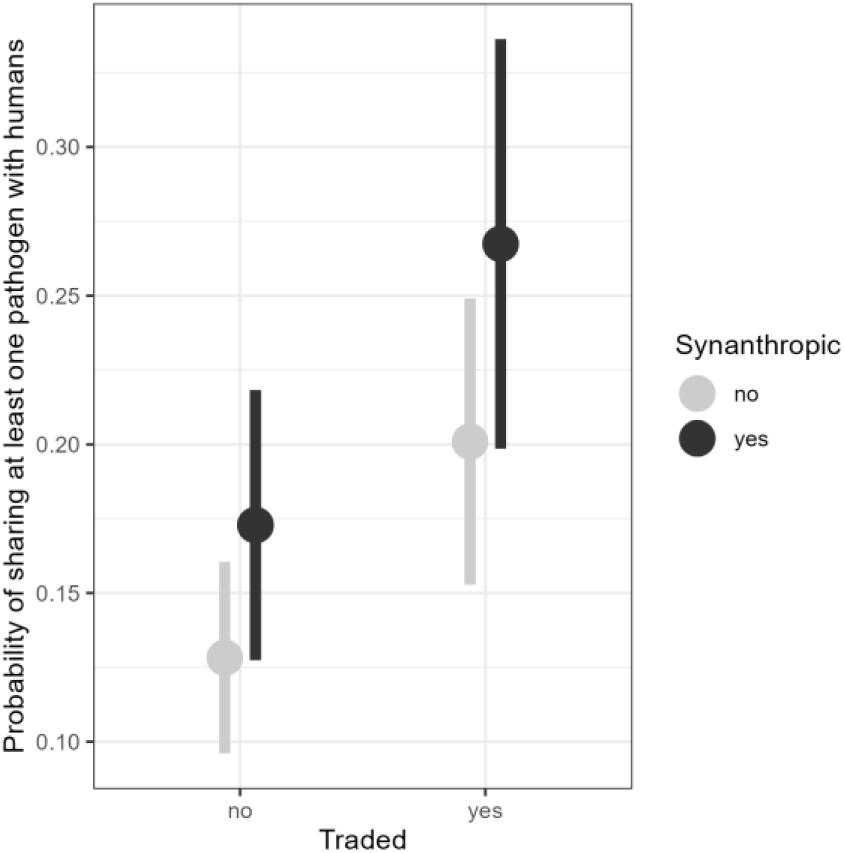
Average effects of trade and synanthropy on the probability of sharing at least one pathogen with humans.

## Supplementary Text 2

Detailed statistics for the Structural Equation Model analysis presented in Fig. 1d.

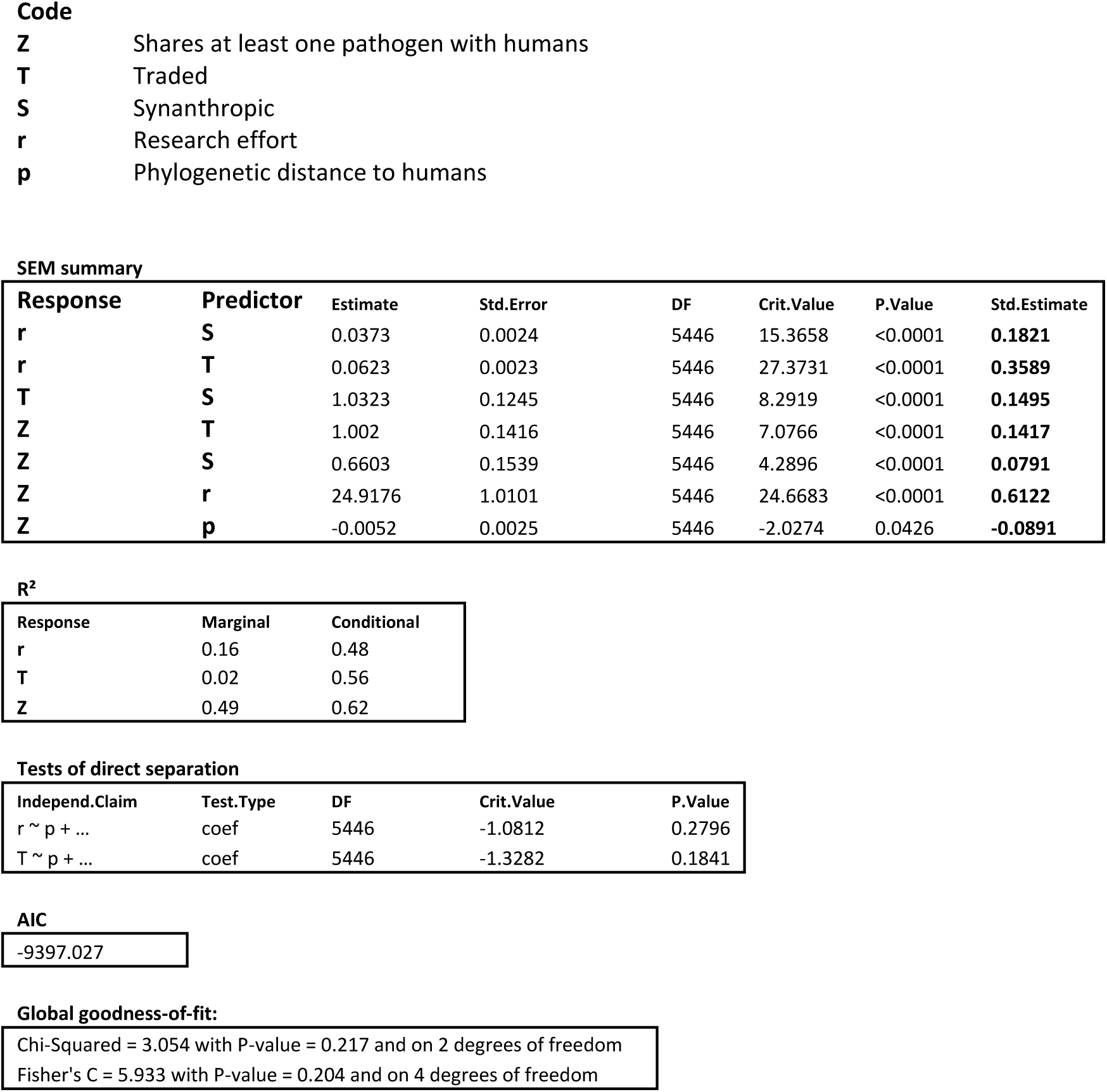

## Supplementary Text 3

Detailed results of the negative binomial mixed effects Hurdle model (presented in Fig. 2b) used to test whether time in trade predicts the probability that wild mammal species share pathogens with humans, and the number of pathogens shared.

| AIC | BIC | logLik | deviance | df.resid |
| --- | --- | --- | --- | --- |
| 1746.3 | 1810.9 | -858.2 | 1716.3 | 531 |

Random effects, zero-inflation
| Groups | Name | Variance | Std.Dev. |
| --- | --- | --- | --- |
| BioRealm | (Intercept) | 0.0444 | 0.2107 |
| family | (Intercept) | 0.3849 | 0.6204 |
| Number of obs: 546, groups: BioRealm, 8; family, 40 |  |  |  |

Random effects, conditional
| Groups | Name | Variance | Std.Dev. |
| --- | --- | --- | --- |
| genus:family | (Intercept) | 0.14713 | 0.3836 |
| family | (Intercept) | 0.07551 | 0.2748 |
| Number of obs: 546, groups: genus:family, 193; family, 40 |  |  |  |

Fixed effects, zero-inflation
|  | Estimate | Std. Error | z value | p-value |  |
| --- | --- | --- | --- | --- | --- |
| (Intercept) | 0.22561 | 0.52778 | 0.427 | 0.669 | n.s. |
| Time in trade | -0.110438 | 0.016393 | -6.737 | <0.0001 | *** |
| Synanthropy [yes] | 0.15854 | 0.452176 | 0.351 | 0.726 | n.s. |
| Research effort | -15.311148 | 2.345604 | -6.528 | <0.0001 | *** |
| Phylo. dist. to humans | 0.025876 | 0.004338 | 5.965 | <0.0001 | *** |

Fixed effects, conditional
|  | Estimate | Std. Error | z value | p-value |  |
| --- | --- | --- | --- | --- | --- |
| (Intercept) | -0.223679 | 0.270244 | -0.828 | 0.408 | n.s. |
| Time in trade | 0.031125 | 0.005669 | 5.49 | <0.0001 | *** |
| Synanthropy [yes] | -0.055035 | 0.186671 | -0.295 | 0.768 | n.s. |
| Research effort | 9.424277 | 1.036089 | 9.096 | <0.0001 | *** |
| Phylo. dist. to humans | -0.008383 | 0.001803 | -4.65 | <0.0001 | *** |

Effects size, zero-inflation
|  | estimate | conf.low | conf.high | statistic | p.value | Effect size |
| --- | --- | --- | --- | --- | --- | --- |
| TradeFrequency | 0.2298062 | 0.1498161 | 0.3525047 | -6.7367423 | <0.0001 | **0.23\*\*\*** |
| Synanthropy [yes] | 1.171799 | 0.4830148 | 2.8427969 | 0.3506163 | 0.726 | **1.17** |
| Research effort | 0.2479337 | 0.1631061 | 0.3768782 | -6.5272465 | <0.0001 | **0.25\*\*\*** |
| Phylo. dist. to humans | 5.2984494 | 3.0630723 | 9.1651662 | 5.9636804 | <0.0001 | **5.3\*\*\*** |

Effects size, conditional
|  | estimate | conf.low | conf.high | statistic | p.value | Effect size |
| --- | --- | --- | --- | --- | --- | --- |
| TradeFrequency | 1.5135212 | 1.3053556 | 1.7548831 | 5.4897721 | <0.0001 | **1.51\*\*\*** |
| Synanthropy [yes] | 0.9464512 | 0.6564525 | 1.3645615 | -0.2948279 | 0.768 | **0.95** |
| Research effort | 2.3593736 | 1.9609093 | 2.8388074 | 9.0947895 | <0.0001 | **2.36\*\*\*** |
| Phylo. dist. to humans | 0.5826235 | 0.4639791 | 0.7316064 | -4.6499482 | <0.0001 | **0.58\*\*** |

## Supplementary Text 4

Interpretable effect sizes for zero-inflation sub-model. Zero-inflation effect sizes correspond to odds ratio for the probability of observing 0. We thus need to calculate the reciprocal to interpret as the probability of observing 1 or more (for example: 1/0.2298062 = 4.3514926; corresponding values highlighted in the following tables).

|  | estimate | conf.low | conf.high |
| --- | --- | --- | --- |
| TradeFrequency | 0.2298062 | 0.1498161 | 0.3525047 |
| Synanthropy [yes] | 1.171799 | 0.4830148 | 2.8427969 |
| Research effort | 0.2479337 | 0.1631061 | 0.3768782 |
| Phylo. dist. to humans | 5.2984494 | 3.0630723 | 9.1651662 |

Interpretable effect sizes.
|  | estimate | conf.low | conf.high |
| --- | --- | --- | --- |
| TradeFrequency | 4.3514926 | 2.8368416 | 6.6748483 |
| Synanthropy [yes] | 0.8533887 | 0.3517663 | 2.0703299 |
| Research effort | 4.0333366 | 2.6533774 | 6.1309802 |
| Phylo. dist. to humans | 0.1887345 | 0.1091088 | 0.3264696 |

## Supplementary Text 5

Estimation of the rate at which species share new pathogens with humans as a function of time in trade.

First, we computed the predicted average response (i.e., number of pathogens shared with humans) across the range of the predictor variable (i.e., time in trade) using *ggaverage* from the *ggeffects* package (Fig. S3a). Average response corresponds to population-level predictions that are representative of covariate distribution.

We then calculated the numerical derivative of this average effect using finite differences, calculated as:

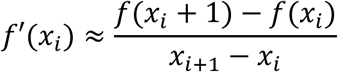

Where 𝑓(𝑥_𝑖_) is the predicted average effect at the 𝑖-th value of the predictor variable. To interpret this rate of change, we calculated its reciprocal (1/𝑓′(𝑥)) which represents the number of years in trade needed to observe an increase of 1 in the response variable: the number of pathogens shared with humans (Fig. S3b). We finally calculated the mean±CI95% rate of change over the whole range of the predictor variable (Fig. S3b).

**Fig. S3:**
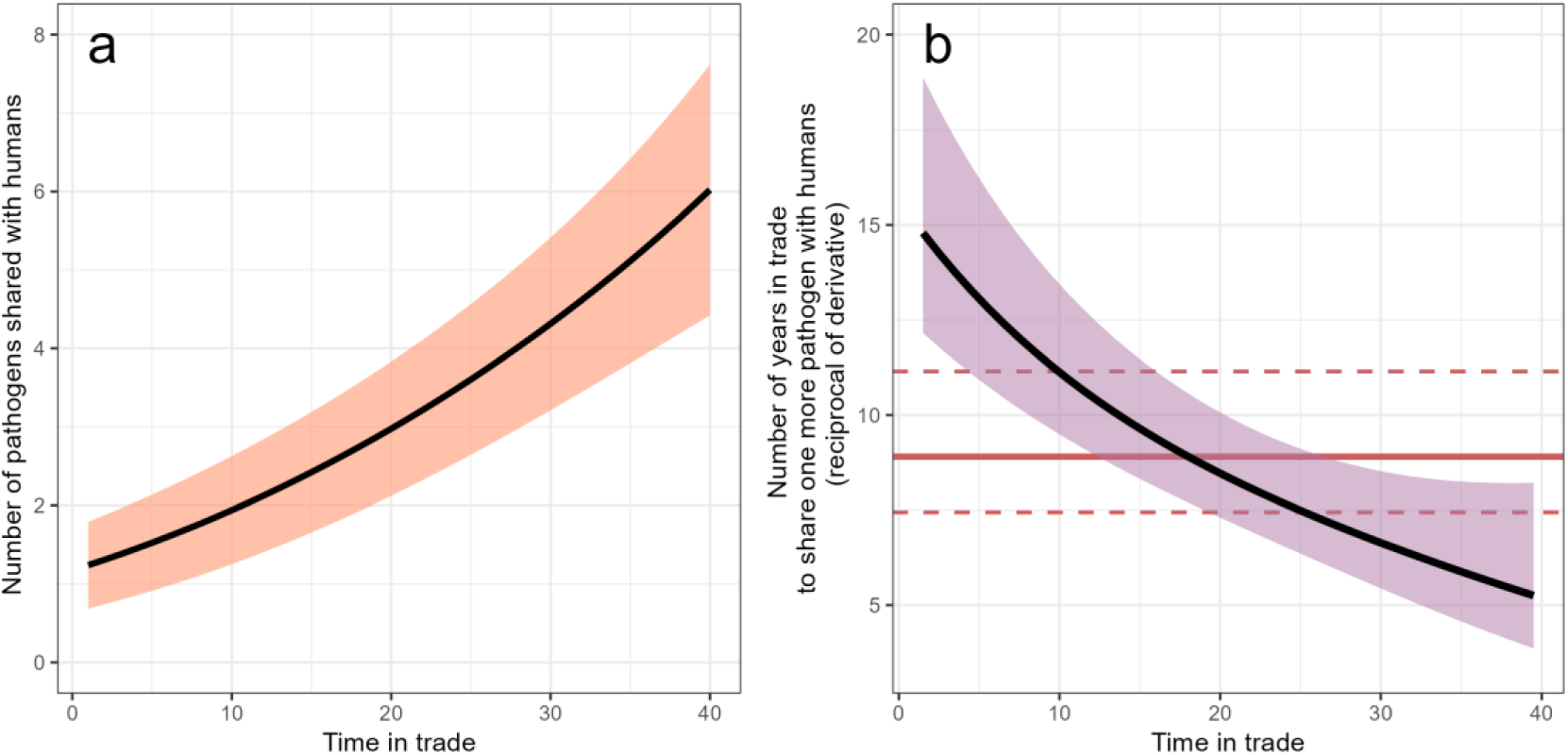
Estimation of the rate at which species share new pathogens with humans as a function of time in trade. **a**, Average response based on the negative binomial mixed effects Hurdle model presented in the article (see Fig. 2), with CI95% around the mean slope. **b**, Reciprocal of the derivative (instantaneous rate of change) of the function presented in panel **a** (with the reciprocals of associated CI95%). This black curve represents the number of years that a species needs to spend in trade to share one more pathogen with human, for a given value of time in trade. The red horizontal solid line represents the average number of years over the whole range of time in trade (8.9 years) and the dotted horizontal red lines the equivalent for the minimum and maximum CI95%, respectively 7.4 and 11.2 years.

## Supplementary Text 6

### Sensitivity analysis

Two aspects of the dataset used to assess host-pathogen associations (CLOVER) were filtered out independently (resulting in four possible combinations, and thus datasets) and analyses were repeated on each dataset to test whether our results were robust or not. The A1 dataset (complete dataset) is the one presented in the main manuscript. All datasets lead to the same conclusions, with only negligible changes in models’ results.

|  |  | EID2 excluded |  |
| --- | --- | --- | --- |
|  |  | No | Yes |
| Viruses only | No | A1 | A2 |
|  | Yes | A3 | A4 |

Detailed description of the filtered datasets:

**A1** – Full dataset with all pathogen types (viruses, bacteria, helminths, fungi, protozoa), includes EID2 data. Dataset used in the results presented in the article.

**A2** – All pathogen types, excludes EID2 data.

**A3** – Viruses only, includes EID2 data.

**A4** – Viruses only, excludes EID2 data.

1) **First analysis (Fig. 1c):** Testing whether traded species are more likely to share pathogen with humans than non-traded species (Binomial mixed effects model)

**Fig. S4:**
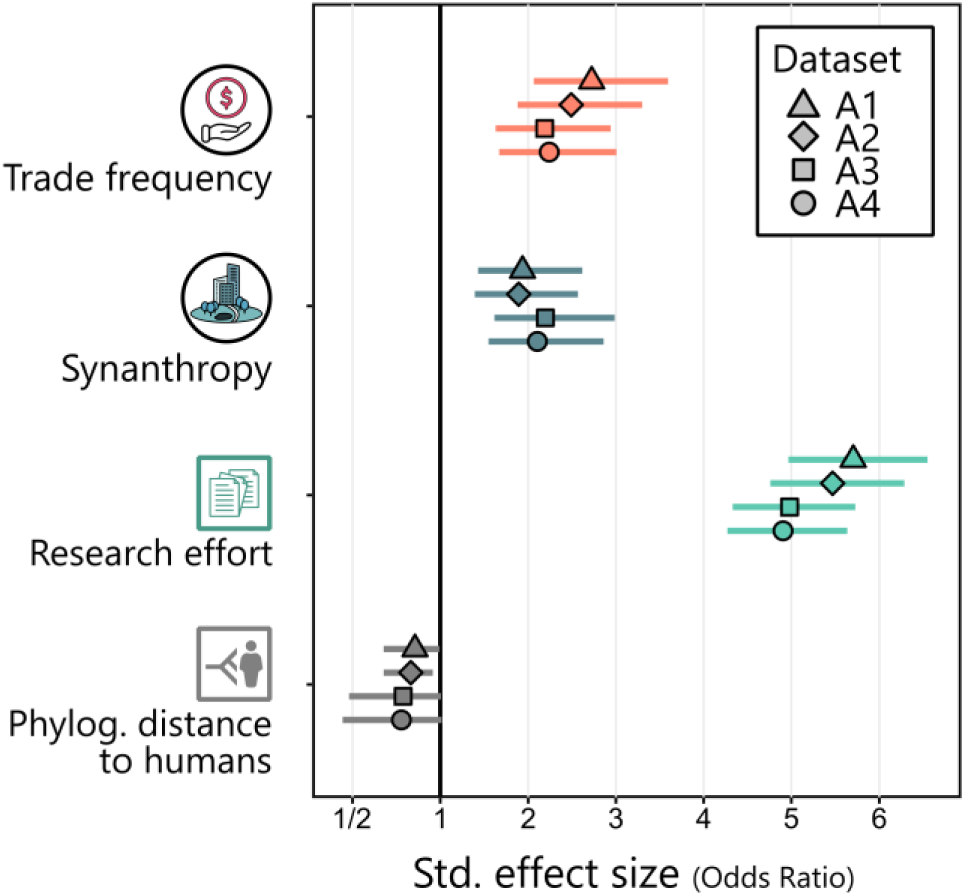
Comparison of predictors’ effect sizes among datasets. All datasets lead to approximately the same results.

Number of species with 1+ zoonotic pathogens: **A1: 871, A2: 799, A3: 699, A4: 691**. Detailed summary of each model:

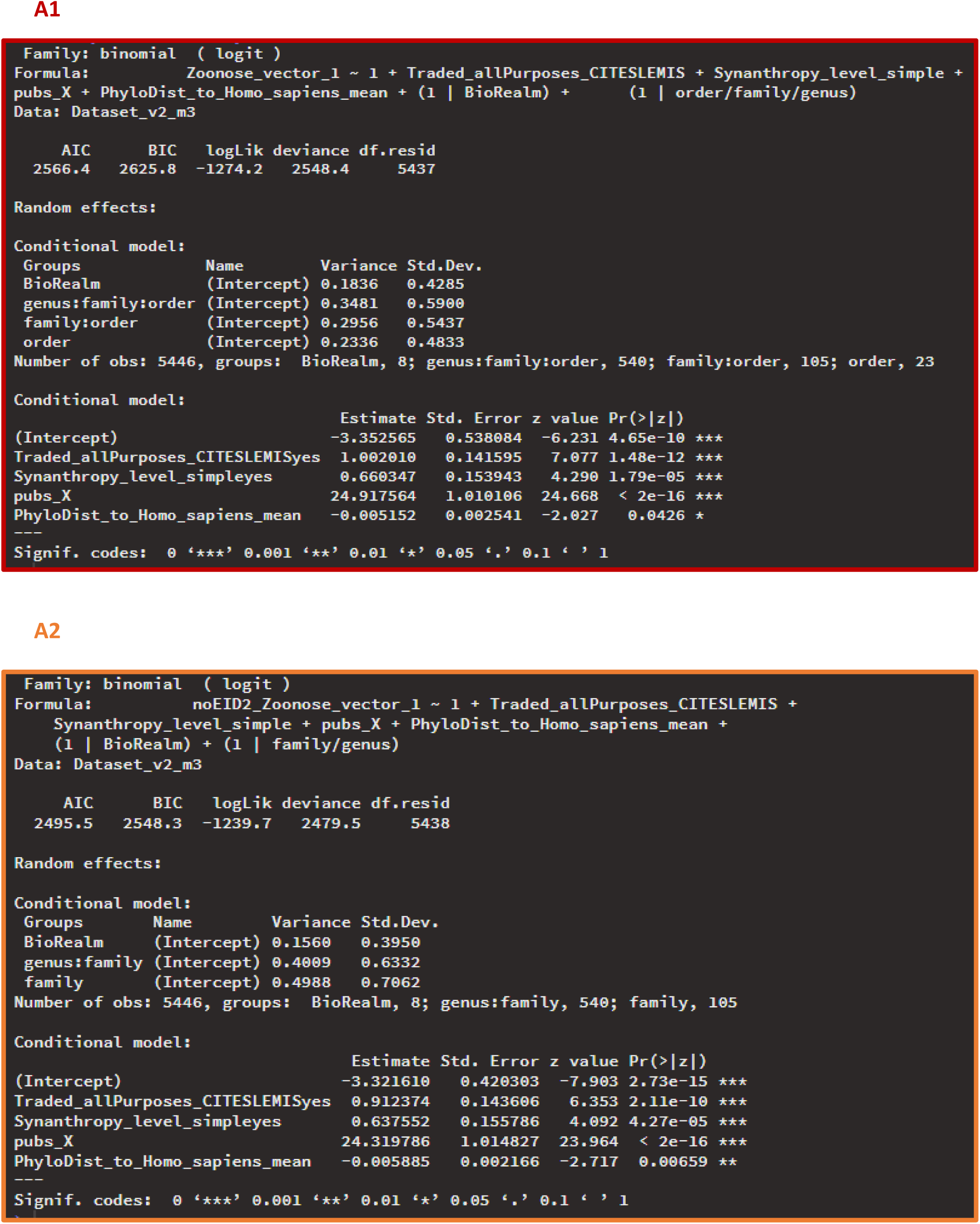
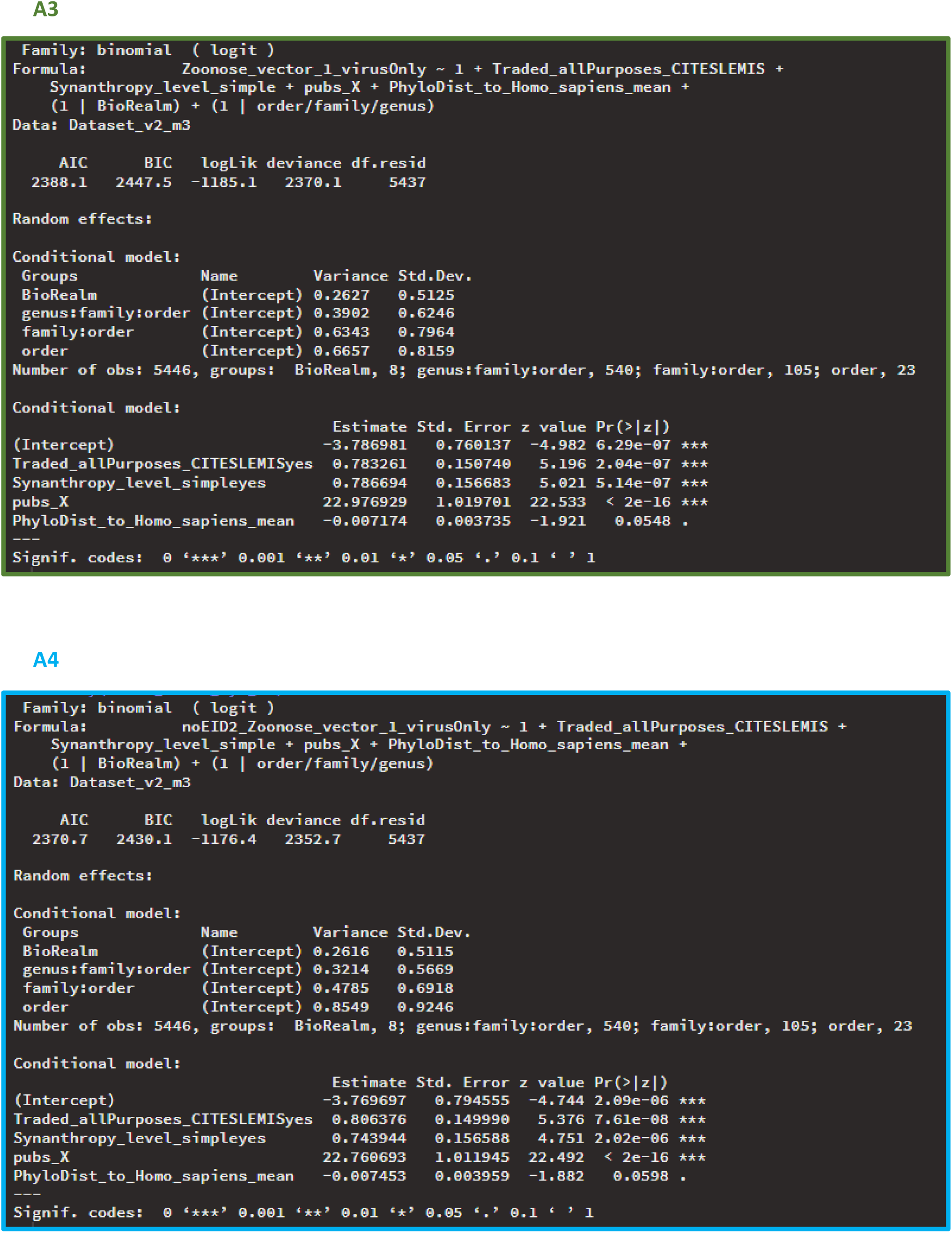

2) **Second analysis (Fig. 1d):** Testing whether traded species are more likely to share pathogen with humans than non-traded species (Structural Equation Model).

We found only minor differences between the 4 datasets. The main one being the non-significant effect of phylogenetic distance to humans in datasets A3 and A4 (p=0.0548 and 0.0598, respectively). Detail summaries of each SEM are displayed hereafter.

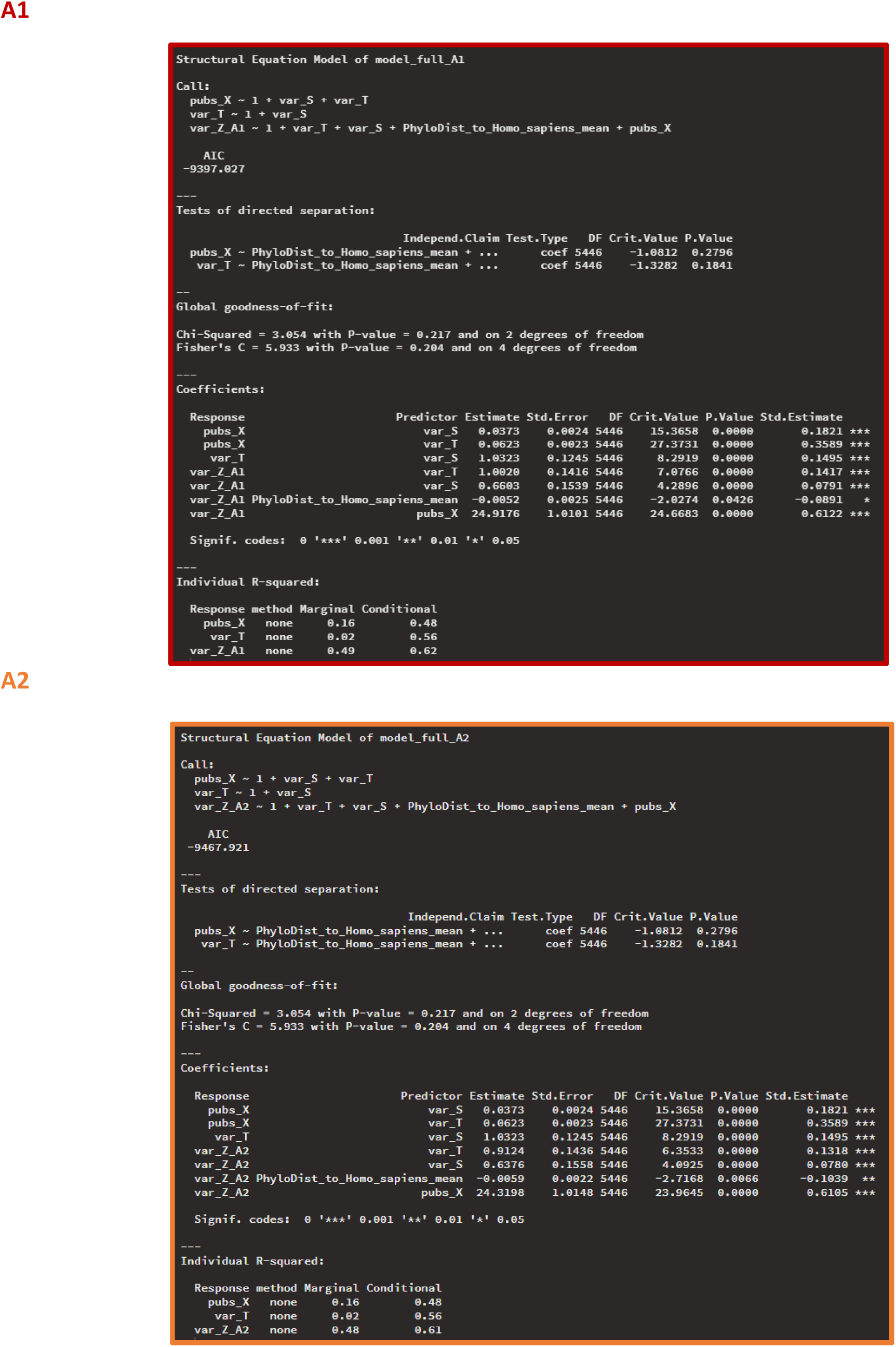
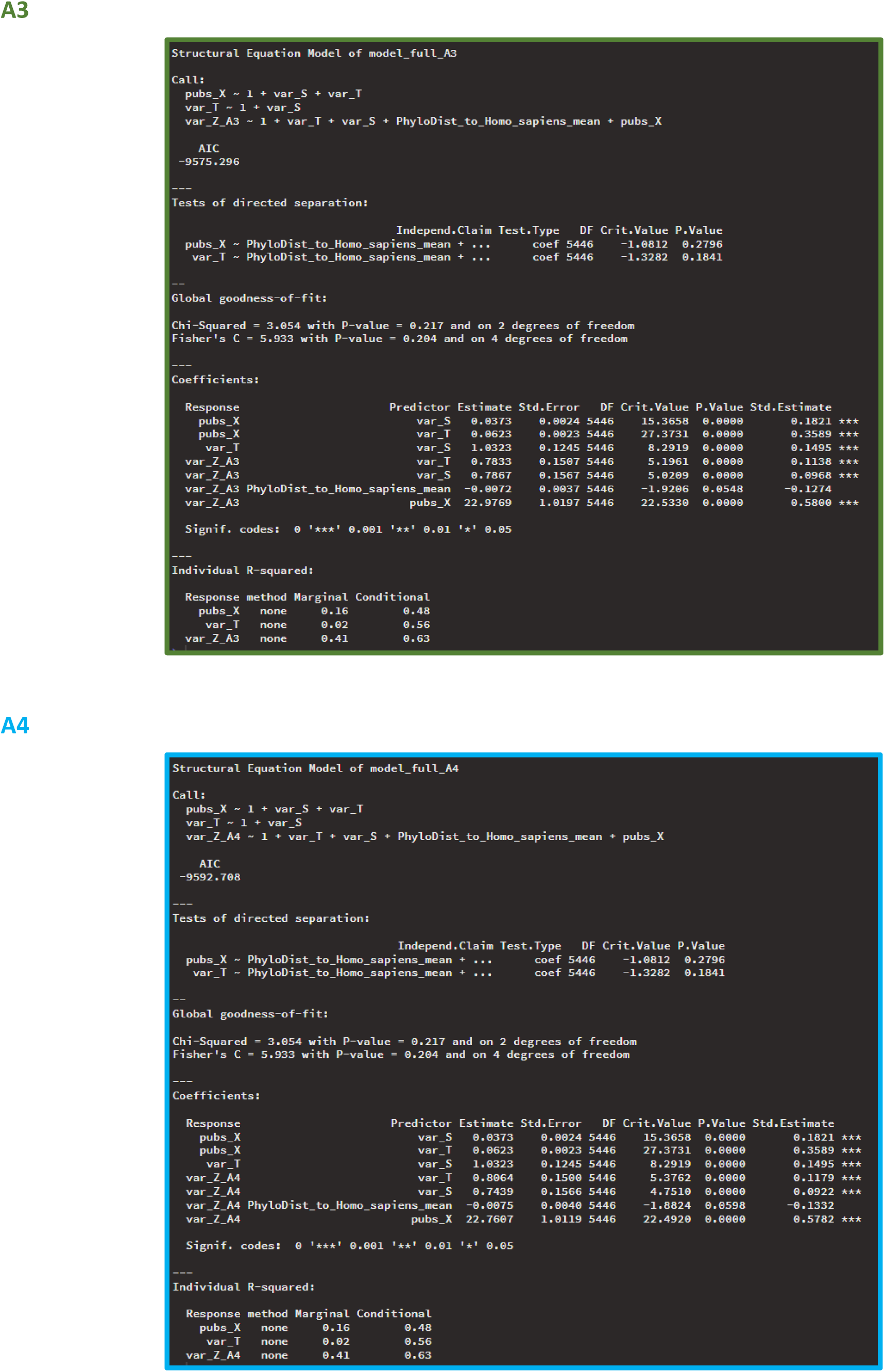

3) **Third analysis (Fig. 2):** Testing whether time in trade (the number of years in which a species has been traded at global scale between 1980 and 2019) is linked to the probability that species share at least one pathogen with humans (zero-inflation sub-model), and the number of pathogens shared with humans (conditional sub-model).

**Fig. S5:**
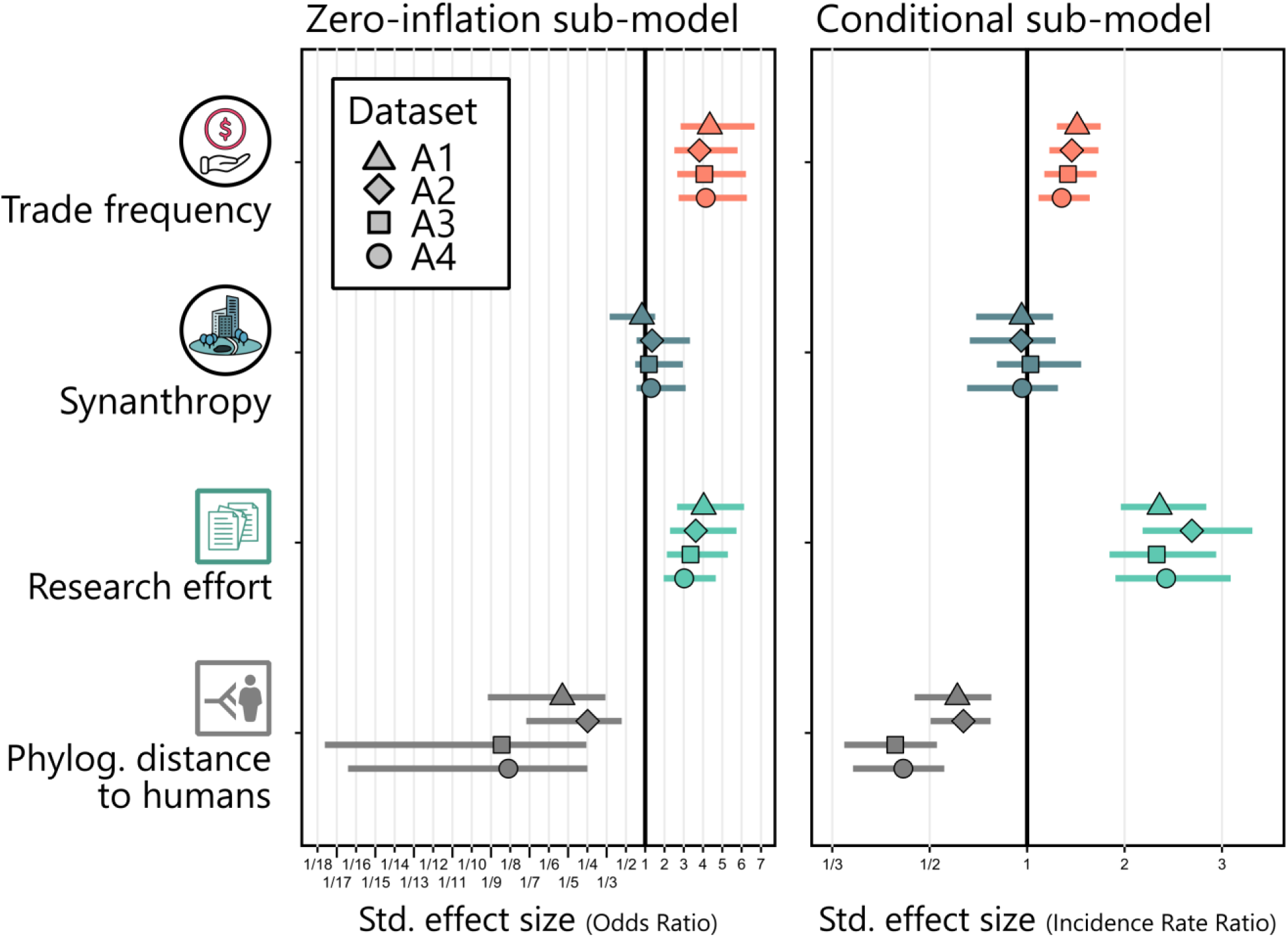
Comparison of predictors’ effect sizes among datasets. All datasets lead to approximately the same results.

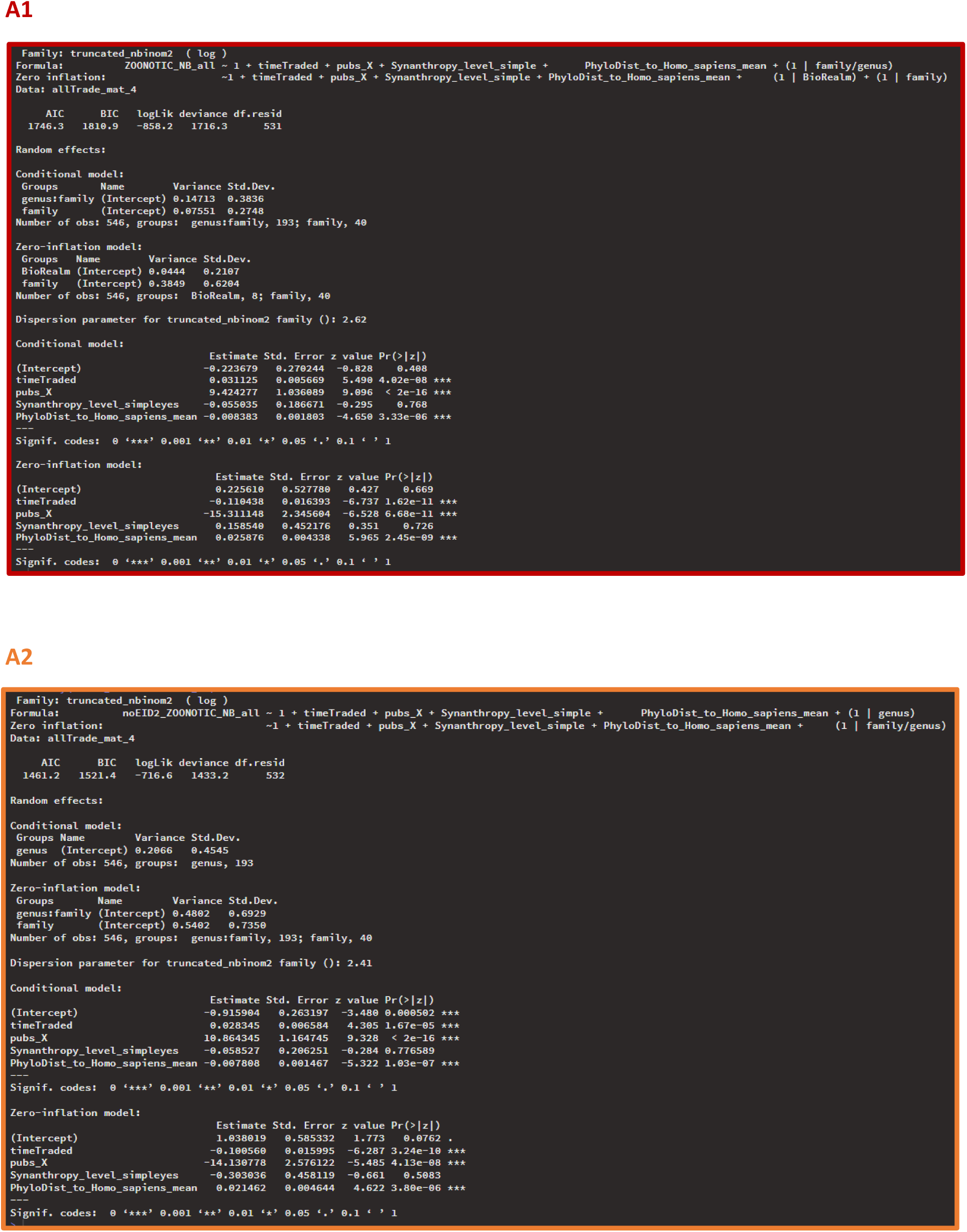
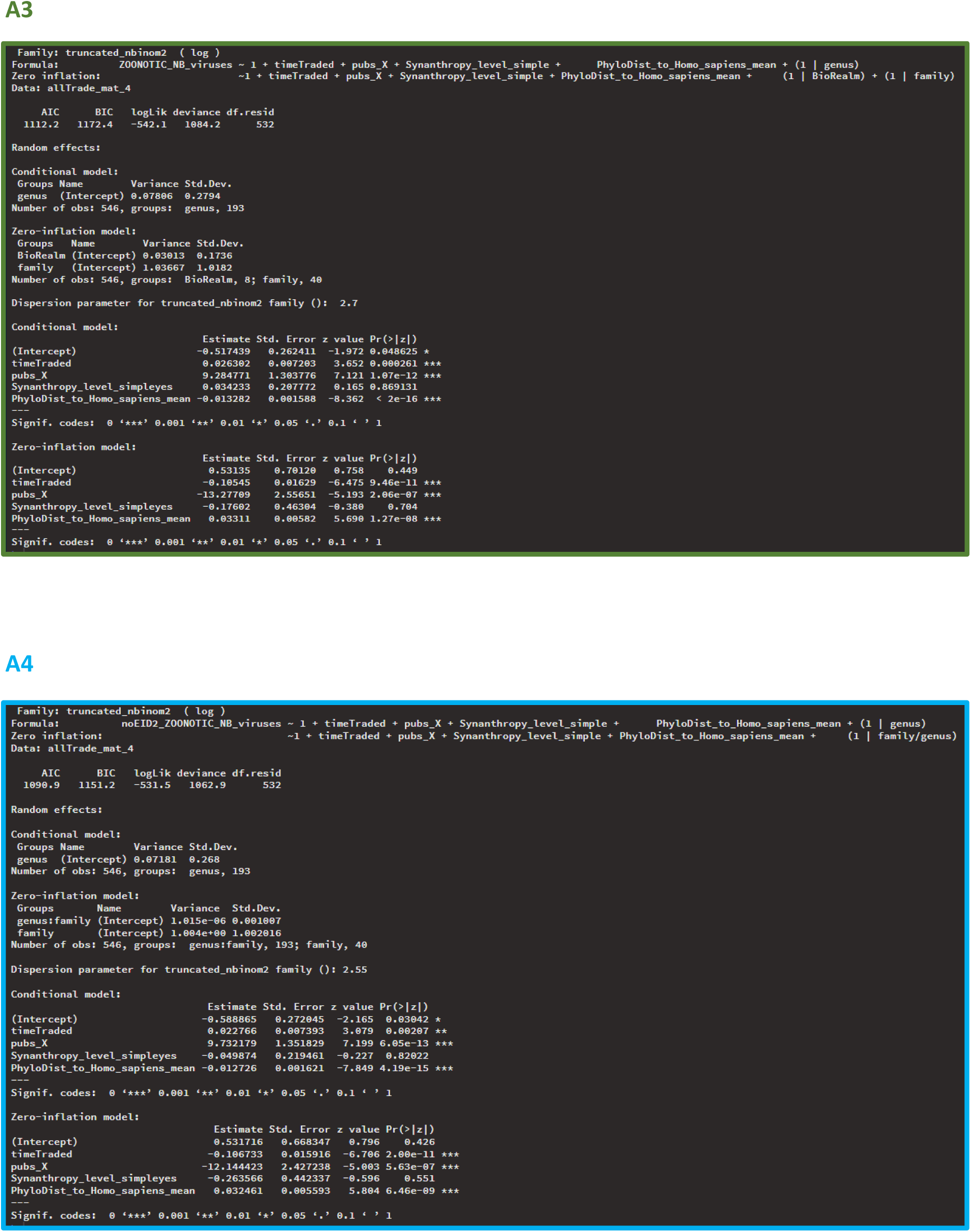

## Notes

### Competing Interest Statement

The authors have declared no competing interest.

## References

1. B. R. Scheffers, B. F. Oliveira, I. Lamb, D. P. Edwards, Global wildlife trade across the tree of life. Science 366, 71–76 (2019).

2. W. B. Karesh, R. A. Cook, E. L. Bennett, J. Newcomb, Wildlife Trade and Global Disease Emergence. Emerg. Infect. Dis. 11, 1000–1002 (2005).

3. B. B. Chomel, A. Belotto, F.-X. Meslin, Wildlife, Exotic Pets, and Emerging Zoonoses. Emerg. Infect. Dis. 13, 6–11 (2007).

4. A. C. Fagre, L. E. Cohen, E. A. Eskew, M. Farrell, E. Glennon, M. B. Joseph, H. K. Frank, S. J. Ryan, C. J. Carlson, G. F. Albery, Assessing the risk of human-to-wildlife pathogen transmission for conservation and public health. Ecology Letters 25, 1534–1549 (2022).

5. S. Friant, Human behaviors driving disease emergence. Evolutionary Anthropology 33, e22015 (2024).

6. K. E. Jones, N. G. Patel, M. A. Levy, A. Storeygard, D. Balk, J. L. Gittleman, P. Daszak, Global trends in emerging infectious diseases. Nature 451, 990–993 (2008).

7. K. J. Olival, P. R. Hosseini, C. Zambrana-Torrelio, N. Ross, T. L. Bogich, P. Daszak, Host and viral traits predict zoonotic spillover from mammals. Nature 546, 646–650 (2017).

8. P. Hosseinzadeh, M. Zareipour, E. Baljani, M. R. Moradali, Social Consequences of the COVID-19 Pandemic. A Systematic Review. invest. educ. enferm 40 (2022).

9. Mortality Analyses, Johns Hopkins Coronavirus Resource Center. https://coronavirus.jhu.edu/data/mortality.

10. T. Allen, K. A. Murray, C. Zambrana-Torrelio, S. S. Morse, C. Rondinini, M. Di Marco, N. Breit, K. J. Olival, P. Daszak, Global hotspots and correlates of emerging zoonotic diseases. Nat Commun 8, 1124 (2017).

11. F. Ecke, B. A. Han, B. Hörnfeldt, H. Khalil, M. Magnusson, N. J. Singh, R. S. Ostfeld, Population fluctuations and synanthropy explain transmission risk in rodent-borne zoonoses. Nat Commun 13, 7532 (2022).

12. A. P. Mendoza, A. Muñoz-Maceda, B. M. Ghersi, M. De La Puente, C. Zariquiey, N. Cavero, Y. Murillo, M. Sebastian, Y. Ibañez, P. G. Parker, A. Perez, M. Uhart, J. Robinson, S. H. Olson, M. H. Rosenbaum, Diversity and prevalence of zoonotic infections at the animal-human interface of primate trafficking in Peru. PLoS ONE 19, e0287893 (2024).

13. P. Nawtaisong, M. T. Robinson, K. Khammavong, P. Milavong, A. Rachlin, S. Dittrich, A. Dubot-Pérès, M. Vongsouvath, P. F. Horwood, P. Dussart, W. Theppangna, B. Douangngeum, A. E. Fine, M. Pruvot, P. N. Newton, Zoonotic Pathogens in Wildlife Traded in Markets for Human Consumption, Laos. Emerg. Infect. Dis. 28, 860–864 (2022).

14. N. Q. Huong, N. T. T. Nga, N. V. Long, B. D. Luu, A. Latinne, M. Pruvot, N. T. Phuong, L. T. V. Quang, V. V. Hung, N. T. Lan, N. T. Hoa, P. Q. Minh, N. T. Diep, N. Tung, V. D. Ky, S. I. Roberton, H. B. Thuy, N. V. Long, M. Gilbert, L. Wicker, J. A. K. Mazet, C. K. Johnson, T. Goldstein, A. Tremeau-Bravard, V. Ontiveros, D. O. Joly, C. Walzer, A. E. Fine, S. H. Olson, Coronavirus testing indicates transmission risk increases along wildlife supply chains for human consumption in Viet Nam, 2013-2014. PLoS ONE 15, e0237129 (2020).

15. L. J. Hughes, O. Morton, B. R. Scheffers, D. P. Edwards, The ecological drivers and consequences of wildlife trade. Biological Reviews 98, 775–791 (2023).

16. M. Worobey, J. I. Levy, L. Malpica Serrano, A. Crits-Christoph, J. E. Pekar, S. A. Goldstein, A. L. Rasmussen, M. U. G. Kraemer, C. Newman, M. P. G. Koopmans, M. A. Suchard, J. O. Wertheim, P. Lemey, D. L. Robertson, R. F. Garry, E. C. Holmes, A. Rambaut, K. G. Andersen, The Huanan Seafood Wholesale Market in Wuhan was the early epicenter of the COVID-19 pandemic. Science 377, 951–959 (2022).

17. P. Lemey, O. G. Pybus, B. Wang, N. K. Saksena, M. Salemi, A.-M. Vandamme, Tracing the origin and history of the HIV-2 epidemic. Proc. Natl. Acad. Sci. U.S.A. 100, 6588–6592 (2003).

18. S. D. Judson, R. Fischer, A. Judson, V. J. Munster, Ecological Contexts of Index Cases and Spillover Events of Different Ebolaviruses. PLoS Pathog 12, e1005780 (2016).

19. J. Cohen, Monkeypox could establish new reservoirs in animals. Science 376, 1258–1259 (2022).

20. K. Paphitis, C. A. Habrun, G. S. Stapleton, A. Reid, C. Lee, A. Majury, A. Murphy, H. McClinchey, A. Corbeil, A. Kearney, K. Benedict, B. Tolar, R. O. Forrest, *Salmonella* Vitkin Outbreak Associated with Bearded Dragons, Canada and United States, 2020–2022. Emerg. Infect. Dis. 30 (2024).

21. K. N. Shivaprakash, S. Sen, S. Paul, J. M. Kiesecker, K. S. Bawa, Mammals, wildlife trade, and the next global pandemic. Current Biology 31, 3671–3677.e3 (2021).

22. M. A. Bezerra-Santos, J. A. Mendoza-Roldan, R. C. A. Thompson, F. Dantas-Torres, D. Otranto, Illegal Wildlife Trade: A Gateway to Zoonotic Infectious Diseases. Trends in Parasitology 37, 181– 184 (2021).

23. N. D. Wolfe, C. P. Dunavan, J. Diamond, Origins of major human infectious diseases. Nature 447, 279–283 (2007).

24. B. I. Pavlin, L. M. Schloegel, P. Daszak, Risk of Importing Zoonotic Diseases through Wildlife Trade, United States. Emerg. Infect. Dis. 15, 1721–1726 (2009).

25. S. Borsky, H. Hennighausen, A. Leiter, K. Williges, CITES and the Zoonotic Disease Content in International Wildlife Trade. Environ Resource Econ 76, 1001–1017 (2020).

26. M. Wardeh, M. S. C. Blagrove, K. J. Sharkey, M. Baylis, Divide-and-conquer: machine-learning integrates mammalian and viral traits with network features to predict virus-mammal associations. Nat Commun 12, 3954 (2021).

27. J. Choo, L. T. P. Nghiem, A. Benítez-López, L. R. Carrasco, Range area and the fast–slow continuum of life history traits predict pathogen richness in wild mammals. Sci Rep 13, 20191 (2023).

28. A. Tonelli, H. Caceres-Escobar, M. S. C. Blagrove, M. Wardeh, M. Di Marco, Identifying life-history patterns along the fast-slow continuum of mammalian viral carriers. R. Soc. Open Sci. 11, 231512 (2024).

29. G. F. Albery, C. J. Carlson, L. E. Cohen, E. A. Eskew, R. Gibb, S. J. Ryan, A. R. Sweeny, D. J. Becker, Urban-adapted mammal species have more known pathogens. Nat Ecol Evol 6, 794–801 (2022).

30. R. Gibb, D. W. Redding, K. Q. Chin, C. A. Donnelly, T. M. Blackburn, T. Newbold, K. E. Jones, Zoonotic host diversity increases in human-dominated ecosystems. Nature 584, 398–402 (2020).

31. N. Mollentze, D. G. Streicker, Viral zoonotic risk is homogenous among taxonomic orders of mammalian and avian reservoir hosts. Proc. Natl. Acad. Sci. U.S.A. 117, 9423–9430 (2020).

32. C. C. S. Tan, L. Van Dorp, F. Balloux, The evolutionary drivers and correlates of viral host jumps. Nat Ecol Evol 8, 960–971 (2024).

33. M. B. Mahon, A. Sack, O. A. Aleuy, C. Barbera, E. Brown, H. Buelow, D. J. Civitello, J. M. Cohen, L. A. De Wit, M. Forstchen, F. W. Halliday, P. Heffernan, S. A. Knutie, A. Korotasz, J. G. Larson, S. L. Rumschlag, E. Selland, A. Shepack, N. Vincent, J. R. Rohr, A meta-analysis on global change drivers and the risk of infectious disease. Nature 629, 830–836 (2024).

34. A. Desvars-Larrive, A. E. Vogl, G. A. Puspitarani, L. Yang, A. Joachim, A. Käsbohrer, A One Health framework for exploring zoonotic interactions demonstrated through a case study. Nat Commun 15, 5650 (2024).

35. J. Choo, L. T. P. Nghiem, S. Chng, L. R. Carrasco, A. Benítez-López, Hotspots of zoonotic disease risk from wildlife hunting and trade in the tropics. Integrative Conservation 2, 165–175 (2023).

36. M. H. Hilderink, I. I. De Winter, No need to beat around the bushmeat–The role of wildlife trade and conservation initiatives in the emergence of zoonotic diseases. Heliyon 7, e07692 (2021).

37. Ö. E. Can, N. D’Cruze, D. W. Macdonald, Dealing in deadly pathogens: Taking stock of the legal trade in live wildlife and potential risks to human health. Global Ecology and Conservation 17, e00515 (2019).

38. R. K. Plowright, C. R. Parrish, H. McCallum, P. J. Hudson, A. I. Ko, A. L. Graham, J. O. Lloyd-Smith, Pathways to zoonotic spillover. Nat Rev Microbiol 15, 502–510 (2017).

39. B. D. Anderson, A. N. Barnes, S. Umar, X. Guo, T. Thongthum, G. C. Gray, “Reverse Zoonotic Transmission (Zooanthroponosis): An Increasing Threat to Animal Health” in Zoonoses: Infections Affecting Humans and Animals, A. Sing, Ed. (Springer International Publishing, Cham, 2023; https://link.springer.com/10.1007/978-3-030-85877-3_59-1), pp. 1–63.

40. K. M. Milich, S. S. Morse, The reverse zoonotic potential of SARS-CoV-2. Heliyon 10, e33040 (2024).

41. CITES trade database, version 2023.1 (2023); https://trade.cites.org.

42. K. M. Smith, C. Zambrana-Torrelio, A. White, M. Asmussen, C. Machalaba, S. Kennedy, K. Lopez, T. M. Wolf, P. Daszak, D. A. Travis, W. B. Karesh, Summarizing US Wildlife Trade with an Eye Toward Assessing the Risk of Infectious Disease Introduction. EcoHealth 14, 29–39 (2017).

43. E. A. Eskew, A. M. White, N. Ross, K. M. Smith, K. F. Smith, J. P. Rodríguez, C. Zambrana-Torrelio, W. B. Karesh, P. Daszak, United States wildlife and wildlife product imports from 2000–2014. Sci Data 7, 22 (2020).

44. CITES Secretariat, “World Wildlife Trade Report” (Geneva, Switzerland, 2022).

45. B. J. Weissgold, US wildlife trade data lack quality control necessary for accurate scientific interpretation and policy application. Conservation Letters. 17, e13005 (2024).

46. A. Etard, S. Morrill, T. Newbold, Global gaps in trait data for terrestrial vertebrates. Global Ecol. Biogeogr. 29, 2143–2158 (2020).

47. IUCN, The IUCN Red List of Threatened Species, IUCN Red List of Threatened Species. https://www.iucnredlist.org/en.

48. R. Gibb, G. F. Abery, D. J. Becker, L. Brierley, R. Connor, T. A. Dallas, E. A. Eskew, M. J. Farrell, A. L. Rasmussen, S. J. Ryan, A. Sweeny, C. J. Carlson, T. Poisot, Data proliferation, reconciliation, and synthesis in viral ecology. BioScience 71, 1148–1156 (2021).

49. K. A. Murray, N. Preston, T. Allen, C. Zambrana-Torrelio, P. R. Hosseini, P. Daszak, Global biogeography of human infectious diseases. Proc. Natl. Acad. Sci. U.S.A. 112, 12746–12751 (2015).

50. A.-S. Stensgaard, R. R. Dunn, B. J. Vennervald, C. Rahbek, The neglected geography of human pathogens and diseases. Nat Ecol Evol 1, 0190 (2017).

51. B. A. Han, A. M. Kramer, J. M. Drake, Global Patterns of Zoonotic Disease in Mammals. Trends in Parasitology 32, 565–577 (2016).

52. R. Gibb, G. F. Albery, N. Mollentze, E. A. Eskew, L. Brierley, S. J. Ryan, S. N. Seifert, C. J. Carlson, Mammal virus diversity estimates are unstable due to accelerating discovery effort. Biolgy Letters (2021).

53. B. A. Betke, N. L. Gottdenker, L. A. Meyers, D. J. Becker, Ecological and evolutionary characteristics of anthropogenic roosting ability in bats of the world. iScience 27, 110369 (2024).

54. D. Liang, X. Giam, S. Hu, L. Ma, D. S. Wilcove, Assessing the illegal hunting of native wildlife in China. Nature 623, 100–105 (2023).

55. M. Harfoot, S. A. M. Glaser, D. P. Tittensor, G. L. Britten, C. McLardy, K. Malsch, N. D. Burgess, Unveiling the patterns and trends in 40 years of global trade in CITES-listed wildlife. Biological Conservation 223, 47–57 (2018).

56. M. T. Bager Olsen, J. Geldmann, M. Harfoot, D. P. Tittensor, B. Price, P. Sinovas, K. Nowak, N. J. Sanders, N. D. Burgess, Thirty-six years of legal and illegal wildlife trade entering the USA. Oryx 55, 432–441 (2021).

57. T. M. Keesey, PhyloPic. https://www.phylopic.org.

58. S. Morand, K. M. McIntyre, M. Baylis, Domesticated animals and human infectious diseases of zoonotic origins: Domestication time matters. *Infection*, Genetics and Evolution 24, 76–81 (2014).

59. E. C. Holmes, The Ecology of Viral Emergence. Annu. Rev. Virol. 9, 173–192 (2022).

60. Y. Kane, G. Wong, G. F. Gao, Animal Models, Zoonotic Reservoirs, and Cross-Species Transmission of Emerging Human-Infecting Coronaviruses. Annu. Rev. Anim. Biosci. 11, 1–31 (2023).

61. J. A. Blanchong, S. J. Robinson, M. D. Samuel, J. T. Foster, Application of genetics and genomics to wildlife epidemiology. J Wildl Manag 80, 593–608 (2016).

62. W.-T. He, X. Hou, J. Zhao, J. Sun, H. He, W. Si, J. Wang, Z. Jiang, Z. Yan, G. Xing, M. Lu, M. A. Suchard, X. Ji, W. Gong, B. He, J. Li, P. Lemey, D. Guo, C. Tu, E. C. Holmes, M. Shi, S. Su, Virome characterization of game animals in China reveals a spectrum of emerging pathogens. Cell 185, 1117–1129.e8 (2022).

63. J. Zhao, W. Wan, K. Yu, P. Lemey, J. H.-O. Pettersson, Y. Bi, M. Lu, X. Li, Z. Chen, M. Zheng, G. Yan, J. Dai, Y. Li, A. Haerheng, N. He, C. Tu, M. A. Suchard, E. C. Holmes, W.-T. He, S. Su, Farmed fur animals harbour viruses with zoonotic spillover potential. Nature 634, 228–233 (2024).

64. P. R. Sichewo, T. M. Hlokwe, E. M. C. Etter, A. L. Michel, Tracing cross species transmission of Mycobacterium bovis at the wildlife/livestock interface in South Africa. BMC Microbiol 20, 49 (2020).

65. S. J. Anthony, C. K. Johnson, D. J. Greig, S. Kramer, X. Che, H. Wells, A. L. Hicks, D. O. Joly, N. D. Wolfe, P. Daszak, W. Karesh, W. I. Lipkin, S. S. Morse, PREDICT Consortium, J. A. K. Mazet, T. Goldstein, Global patterns in coronavirus diversity. Virus Evolution 3 (2017).

66. K. M. Smith, S. J. Anthony, W. M. Switzer, J. H. Epstein, T. Seimon, H. Jia, M. D. Sanchez, T. T. Huynh, G. G. Galland, S. E. Shapiro, J. M. Sleeman, D. McAloose, M. Stuchin, G. Amato, S.-O. Kolokotronis, W. I. Lipkin, W. B. Karesh, P. Daszak, N. Marano, Zoonotic Viruses Associated with Illegally Imported Wildlife Products. PLoS ONE 7, e29505 (2012).

67. Y. Feng, G. Kuang, Y. Pan, J. Wang, W. Yang, W. Wu, H. Pan, J. Wang, X. Han, L. Yang, G. Xin, Y. Shan, Q. Gou, X. Liu, D. Guo, G. Liang, E. C. Holmes, Z. Gao, M. Shi, Small mammals in biodiversity hotspot harbor viruses of epidemic potential. [Preprint] (2024). 10.1101/2024.09.10.612213.

68. C. Lalueza-Fox, Museomics. Current Biology 32, R1214–R1215 (2022).

69. M. Hämmerle, M. Guellil, L. Trgovec-Greif, O. Cheronet, S. Sawyer, I. Ruiz-Gartzia, E. Lizano, A. Rymbekova, P. Gelabert, P. Bernardi, S. Han, T. Rattei, V. J. Schuenemann, T. Marques-Bonet, K. Guschanski, S. Calvignac-Spencer, R. Pinhasi, M. Kuhlwilm, Screening great ape museum specimens for DNA viruses. Sci Rep 14, 29806 (2024).

70. M. Hämmerle, A. Rymbekova, P. Gelabert, S. Sawyer, O. Cheronet, P. Bernardi, S. Calvignac-Spencer, M. Kuhlwilm, M. Guellil, R. Pinhasi, Link between Monkeypox Virus Genomes from Museum Specimens and 1965 Zoo Outbreak. Emerg. Infect. Dis. 30 (2024).

71. D. DiEuliis, K. R. Johnson, S. S. Morse, D. E. Schindel, Specimen collections should have a much bigger role in infectious disease research and response. Proc. Natl. Acad. Sci. U.S.A. 113, 4–7 (2016).

72. K. N. Balasubramaniam, P. R. Marty, S. Samartino, A. Sobrino, T. Gill, M. Ismail, R. Saha, B. A. Beisner, S. S. K. Kaburu, E. Bliss-Moreau, M. E. Arlet, N. Ruppert, A. Ismail, S. A. M. Sah, L. Mohan, S. K. Rattan, U. Kodandaramaiah, B. McCowan, Impact of individual demographic and social factors on human–wildlife interactions: a comparative study of three macaque species. Sci Rep 10, 21991 (2020).

73. E. A. Eskew, C. J. Carlson, Overselling wildlife trade bans will not bolster conservation or pandemic preparedness. The Lancet Planetary Health 4, e215–e216 (2020).

74. C. S. Peros, R. Dasgupta, P. Kumar, B. A. Johnson, Bushmeat, wet markets, and the risks of pandemics: Exploring the nexus through systematic review of scientific disclosures. Environmental Science & Policy 124, 1–11 (2021).

75. X. Xiao, C. Newman, C. D. Buesching, D. W. Macdonald, Z.-M. Zhou, Animal sales from Wuhan wet markets immediately prior to the COVID-19 pandemic. Sci Rep 11, 11898 (2021).

76. P. Romero-Vidal, G. Blanco, J. M. Barbosa, M. Carrete, F. Hiraldo, E. C. Pacífico, A. Rojas, A. O. Bermúdez-Cavero, J. A. Díaz-Luque, R. León-Pérez, J. L. Tella, The widespread keeping of wild pets in the Neotropics: An overlooked risk for human, livestock and wildlife health. People and Nature 6, 1023–1035 (2024).

77. Bushmeat Database; https://www2.cifor.org/bushmeat/resources/bushmeat-database/.

78. A. P. Mendoza, S. Shanee, N. Cavero, C. Lujan-Vega, Y. Ibañez, C. Rynaby, M. Villena, Y. Murillo, S. H. Olson, A. Perez, P. G. Parker, M. M. Uhart, D. J. Brightsmith, Domestic networks contribute to the diversity and composition of live wildlife trafficked in urban markets in Peru. Global Ecology and Conservation 37, e02161 (2022).

79. A. Borzée, J. McNeely, K. Magellan, J. R. B. Miller, L. Porter, T. Dutta, K. P. Kadinjappalli, S. Sharma, G. Shahabuddin, F. Aprilinayati, G. E. Ryan, A. Hughes, A. H. Abd Mutalib, A. Z. A. Wahab, D. Bista, S. A. Chavanich, J. L. Chong, G. A. Gale, H. Ghaffari, Y. Ghimirey, V. K. Jayaraj, A. P. Khatiwada, M. Khatiwada, M. Krishna, N. Lwin, P. K. Paudel, C. Sadykova, T. Savini, B. B. Shrestha, C. T. Strine, M. Sutthacheep, E. P. Wong, T. Yeemin, N. Z. Zahirudin, L. Zhang, COVID-19 Highlights the Need for More Effective Wildlife Trade Legislation. Trends in Ecology & Evolution 35, 1052–1055 (2020).

80. The CITES species | CITES. https://cites.org/eng/disc/species.php.

81. Role of CITES in reducing risk of future zoonotic disease emergence associated with international trade | CITES. https://cites.org/eng/topics/role_of_CITES_in_reducing_risk_of_future_zoonotic_disease.

82. Reducing the risk of zoonotic disease emergence associated with international wildlife trade: CITES and WOAH enter new agreement | CITES. https://cites.org/eng/news/cites-woah-memorandum-of-understanding-2024-zoonotic-disease-emergence.

83. The global agreement on pandemics in a nutshell, Consilium. https://www.consilium.europa.eu/en/infographics/towards-an-international-treaty-on-pandemics/.

## Cited literature

1. Mammal Diversity Database, version 1.12.1 (2023); 10.5281/zenodo.10595931.

2. S. A. Chamberlain, E. Szöcs, taxize: taxonomic search and retrieval in R. F1000 Research, 1–30 (2013).

3. F. T. Burbrink, B. I. Crother, C. M. Murray, B. T. Smith, S. Ruane, E. A. Myers, R. A. Pyron, Empirical and philosophical problems with the subspecies rank. Ecology and Evolution 12, e9069 (2022).

4. R. Gibb, G. F. Abery, D. J. Becker, L. Brierley, R. Connor, T. A. Dallas, E. A. Eskew, M. J. Farrell, A. L. Rasmussen, S. J. Ryan, A. Sweeny, C. J. Carlson, T. Poisot, Data proliferation, reconciliation, and synthesis in viral ecology. BioScience 71, 1148–1156 (2021).

5. P. R. Stephens, P. Pappalardo, S. Huang, J. E. Byers, M. J. Farrell, A. Gehman, R. R. Ghai, S. E. Haas, B. Han, A. W. Park, J. P. Schmidt, S. Altizer, V. O. Ezenwa, C. L. Nunn, Global Mammal Parasite Database version 2.0. Ecology 98, 1476–1476 (2017).

6. M. Wardeh, C. Risley, M. K. McIntyre, C. Setzkorn, M. Baylis, Database of host-pathogen and related species interactions, and their global distribution. Sci Data 2, 150049 (2015).

8. L. P. Shaw, A. D. Wang, D. Dylus, M. Meier, G. Pogacnik, C. Dessimoz, F. Balloux, The phylogenetic range of bacterial and viral pathogens of vertebrates. Molecular Ecology 29, 3361–3379 (2020).

9. G. F. Albery, C. J. Carlson, L. E. Cohen, E. A. Eskew, R. Gibb, S. J. Ryan, A. R. Sweeny, D. J. Becker, Urban-adapted mammal species have more known pathogens. Nat Ecol Evol 6, 794–801 (2022).

10. R. Gibb, D. W. Redding, K. Q. Chin, C. A. Donnelly, T. M. Blackburn, T. Newbold, K. E. Jones, Zoonotic host diversity increases in human-dominated ecosystems. Nature 584, 398–402 (2020).

11. L. Zhang, J. Rohr, R. Cui, Y. Xin, L. Han, X. Yang, S. Gu, Y. Du, J. Liang, X. Wang, Z. Wu, Q. Hao, X. Liu, Biological invasions facilitate zoonotic disease emergences. Nat Commun 13, 1762 (2022).

12. D. Fantini, easyPubMed: Search and Retrieve Scientific Publication Records from PubMed, version 2.13 (2019); https://CRAN.R-project.org/package=easyPubMed.

13. PubMed, *PubMed*. https://pubmed.ncbi.nlm.nih.gov/.

14. N. S. Upham, J. A. Esselstyn, W. Jetz, Inferring the mammal tree: Species-level sets of phylogenies for questions in ecology, evolution, and conservation. PLoS Biol 17, e3000494 (2019).

15. IUCN, The IUCN Red List of Threatened Species, IUCN Red List of Threatened Species. https://www.iucnredlist.org/en.

16. CITES trade database, version 2023.1 (2023); https://trade.cites.org.

17. E. A. Eskew, A. M. White, N. Ross, K. M. Smith, K. F. Smith, J. P. Rodríguez, C. Zambrana-Torrelio, W. B. Karesh, P. Daszak, United States wildlife and wildlife product imports from 2000–2014. Sci Data 7, 22 (2020).

18. D. W. S. Challender, D. Brockington, A. Hinsley, M. Hoffmann, J. E. Kolby, F. Massé, D. J. D. Natusch, T. E. E. Oldfield, W. Outhwaite, M. ’T Sas-Rolfes, E. J. Milner-Gulland, Mischaracterizing wildlife trade and its impacts may mislead policy processes. CONSERVATION LETTERS 15, e12832 (2022).

19. CITES Secretariat, “World Wildlife Trade Report” (Geneva, Switzerland, 2022).

20. K. M. Smith, C. Zambrana-Torrelio, A. White, M. Asmussen, C. Machalaba, S. Kennedy, K. Lopez, T. M. Wolf, P. Daszak, D. A. Travis, W. B. Karesh, Summarizing US Wildlife Trade with an Eye Toward Assessing the Risk of Infectious Disease Introduction. EcoHealth 14, 29–39 (2017).

21. B. J. Weissgold, US wildlife trade data lack quality control necessary for accurate scientific interpretation and policy application. Conservation Letters. 17, e13005 (2024).

22. Checklist of CITES species. https://checklist.cites.org/#/en.

23. The CITES species | CITES. https://cites.org/eng/disc/species.php.

24. R core team, R v.4.2.1: A Language and Environment for Statistical Computing, R Foundation for Statistical Computing (2022).

25. Posit team, RStudio: Integrated Development Environment for R, Posit Software (2023); http://www.posit.co/.

26. G. Yu, D. K. Smith, H. Zhu, Y. Guan, T. T. Lam, ggtree: a package for visualization and annotation of phylogenetic trees with their covariates and other associated data. Methods Ecol Evol 8, 28–36 (2017).

27. L.-G. Wang, T. T.-Y. Lam, S. Xu, Z. Dai, L. Zhou, T. Feng, P. Guo, C. W. Dunn, B. R. Jones, T. Bradley, H. Zhu, Y. Guan, Y. Jiang, G. Yu, Treeio: An R Package for Phylogenetic Tree Input and Output with Richly Annotated and Associated Data. Molecular Biology and Evolution 37, 599–603 (2020).

28. H. Wickham, ggplot2: Elegant Graphics for Data Analysis, Springer-Verlag (2016); https://ggplot2.tidyverse.org.

29. M. Brooks E., K. Kristensen, K. Benthem J., van, A. Magnusson, C. Berg W., A. Nielsen, H. Skaug J., M. Mächler, B. Bolker M., glmmTMB Balances Speed and Flexibility Among Packages for Zero-inflated Generalized Linear Mixed Modeling. The R Journal 9, 378 (2017).

30. D. Lüdecke, M. Ben-Shachar, I. Patil, P. Waggoner, D. Makowski, performance: An R Package for Assessment, Comparison and Testing of Statistical Models. JOSS 6, 3139 (2021).

31. F. Hartig, DHARMa: Residual Diagnostics for Hierarchical (Multi-Level / Mixed) Regression Models, version 0.4.6 (2022); https://CRAN.R-project.org/package=DHARMa.

32. D. J. Barr, R. Levy, C. Scheepers, H. J. Tily, Random effects structure for confirmatory hypothesis testing: Keep it maximal. Journal of Memory and Language 68, 255–278 (2013).

33. D. Lüdecke, sjPlot: Data Visualization for Statistics in Social Science, version 2.8.12 (2022); https://CRAN.R-project.org/package=sjPlot.

34. J. S. Lefcheck, PIECEWISESEM : Piecewise structural equation modelling in R for ecology, evolution, and systematics. Methods Ecol Evol 7, 573–579 (2016).

35. C. J. Carlson, C. B. Brookson, C. A. Cummings, R. Gibb, F. W. Halliday, A. M. Heckley, Z. Y. X. Huang, T. Lavelle, H. Robertson, A. Vincente-Santos, C. M. Weets, T. Poisot, Pathogens and planetary change. Nature Reviews Biodiversity, doi: 10.1038/s44358-024-00005-w.

